# Inactivation of a protein hydroxylase complex impairs replication fork restart in cancer and neurodevelopmental disorders

**DOI:** 10.1101/2025.11.22.689938

**Authors:** Tristan J Kennedy, Sally C Fletcher, Uncaar Boora, Erin Fraser-Williams, Arashpreet Kaur, Gareth W Hughes, Eline Hendrix, Joanna R Morris, Stephen J Smerdon, Mathew L Coleman

## Abstract

Protein hydroxylation is a post-translational modification that is commonly catalysed by enzymes of the oxygen- and 2-oxoglutarate (2OG)-dependent oxygenase family. The Jumonji-C (JmjC)-only sub-family of 2OG-oxygenases catalyse the hydroxylation of protein and tRNA substrates involved in fundamental cellular processes. Jumonji-domain 5 (JMJD5) is a highly conserved and essential 2OG-oxygenase that, thus far, is the only arginyl hydroxylase assigned in eukaryotes. We recently reported that JMJD5 hydroxylase activity is required for DNA replication fidelity, and that pathogenic variants cause replication stress (RS) and genome instability (GIN) in a novel neurodevelopmental disorder. Because of the prevalence of RS and GIN in cancer, and reported roles of JMJD5 in tumorigenesis, we here investigate the impact of JMJD5 cancer mutations on its role in replication fidelity. We describe the structural impact of cancer missense mutations on the hydroxylase activity of JMJD5 and its interaction with ‘RCCD1’, an abundant binding partner encoded by a gene associated with susceptibility to a variety of tumour types. We show that the JMJD5:RCCD1 interaction is disrupted by cancer mutations and that the complex is essential for suppressing RS and GIN in tumour cells. Finally, we describe a novel interaction of the complex with RAD51 paralogues and demonstrate the importance of the JMJD5:RCCD1 interaction for normal replication fork restart. Our findings further highlight the importance of protein hydroxylases in fundamental cellular processes and the consequences of JMJD5 and RCCD1 deregulation in human disease.

## Introduction

Mechanistic insight into the basis of disease often follows the characterisation of novel post-translational modifications (PTMs). First reported as a fibrillar collagen modification, protein hydroxylation is the enzymatic incorporation of a single oxygen atom into an amino acid sidechain [1, 2]. Although, the essential role of protein hydroxylation as an oxygen sensor in response to hypoxia has linked the modification intrinsically with disease, it remains poorly characterised compared to other PTMs [3].

The enzymatic family responsible for catalysis of most protein hydroxylation are known as 2OG-oxygenases. Named for their reliance on molecular oxygen, Fe(II) and the Krebs cycle intermediate 2OG as co-factors, these enzymes have been shown to catalyse a range of oxidative modifications on proteins, nucleic acids and lipids [4–6]. This diversity in substrate targets has involved 2OG-oxygenases in many facets of cellular biology. One fundamental pathway involving oxygenation is gene expression, with 2OG-oxygenases identified that control transcription, splicing and translation.

The epigenetic modification of histone tails plays a key role in regulating gene transcription and chromatin structure [7]. The lysine histone demethylases (KDMs) belong to a subfamily of phylogenetically distinct 2OG-oxygenases called Jumonji-C (JmjC) enzymes, which are named after their characteristic catalytic JmjC domain [8]. Epigenetic regulation by KDMs, and its link to human development and disease, has been extensively characterised. However, other JmjC oxygenases with links to gene expression and disease require more comprehensive investigation [5, 9].

JmjC domain protein 5 (JMJD5) is an enigmatic 2OG-oxygenase that is essential for life [10]. JMJD5 belongs to the ‘JmjC-only’ sub-family which, although sharing similarities to KDMs in their catalytic domains, catalyse stable protein hydroxylation rather than demethylation [8]. JMJD5 was originally reported as a KDM, but more recently has been shown to catalyse stable arginyl hydroxylation *in vitro* [11, 12]. We have recently identified human JMJD5 variants that are deleterious to splicing and hydroxylase stability and activity, and that cause a developmental disorder associated with growth delay, intellectual disability, and facial dysmorphism [13]. This clinical phenotype was associated with an increase in RS, which is now recognised as major driver of GIN, including in tumour cells [14]. Although JMJD5 has been linked to cancer [15–19], its importance for replication fidelity and genome stability in this context is unclear. Whether JMJD5 is mutationally inactivated in tumour cells, and the role of this in cancer-associated RS and GIN, is not known.

Here we investigate the role of JMJD5 in replication fidelity using tumour cell models. We describe how JMJD5 is mutated and functionally impaired in a variety of cancers, and that these variants fail to suppress RS. Tumour variants also prevent the interaction of JMJD5 with a novel cancer susceptibility gene product called RCCD1. By discovering interactors of this hydroxylase complex, we assign the function of JMJD5 and RCCD1 to replication fork re-start in both cancer and primary cells derived from neurodevelopmental disorder patients. Our findings provide critical new insights into protein hydroxylase biology, and a solid foundation for mechanistic and preclinical studies of the JMJD5:RCCD1 pathway.

## Results

### JMJD5 knockdown increases basal and stress-induced replication stress in tumour cell models

Our previous work uncovered a role for JMJD5 activity in replication fidelity using patient-derived dermal fibroblasts as a model [13]. To begin to explore the replication role of JMJD5 in other cell types and in cancer, we tested whether JMJD5 controls replication fidelity in four commonly used tumour cell lines; lung and cervical adenocarcinoma epithelial cells (A549 and HeLa, respectively) and bone osteosarcoma cells (U2OS and Saos-2). Following JMJD5 knockdown (Figure 1A and Supplemental Figure 1A/D/G), we quantified micronuclei and 53BP1 bodies as indirect markers of replication stress. Increased micronuclei (Figure 1B, upper panel) were observed in all four models following treatment with two independent JMJD5 siRNAs (Figure 1B, lower panel; Supplemental Figure 1B/E/H). Likewise, we observed increased 53BP1 bodies (Figure 1C, upper panel) following JMJD5 knockdown in each of the cancer cell lines (Figure 1C, lower panel, and Supplemental Figure 1C/F/I).

**Figure 1.**
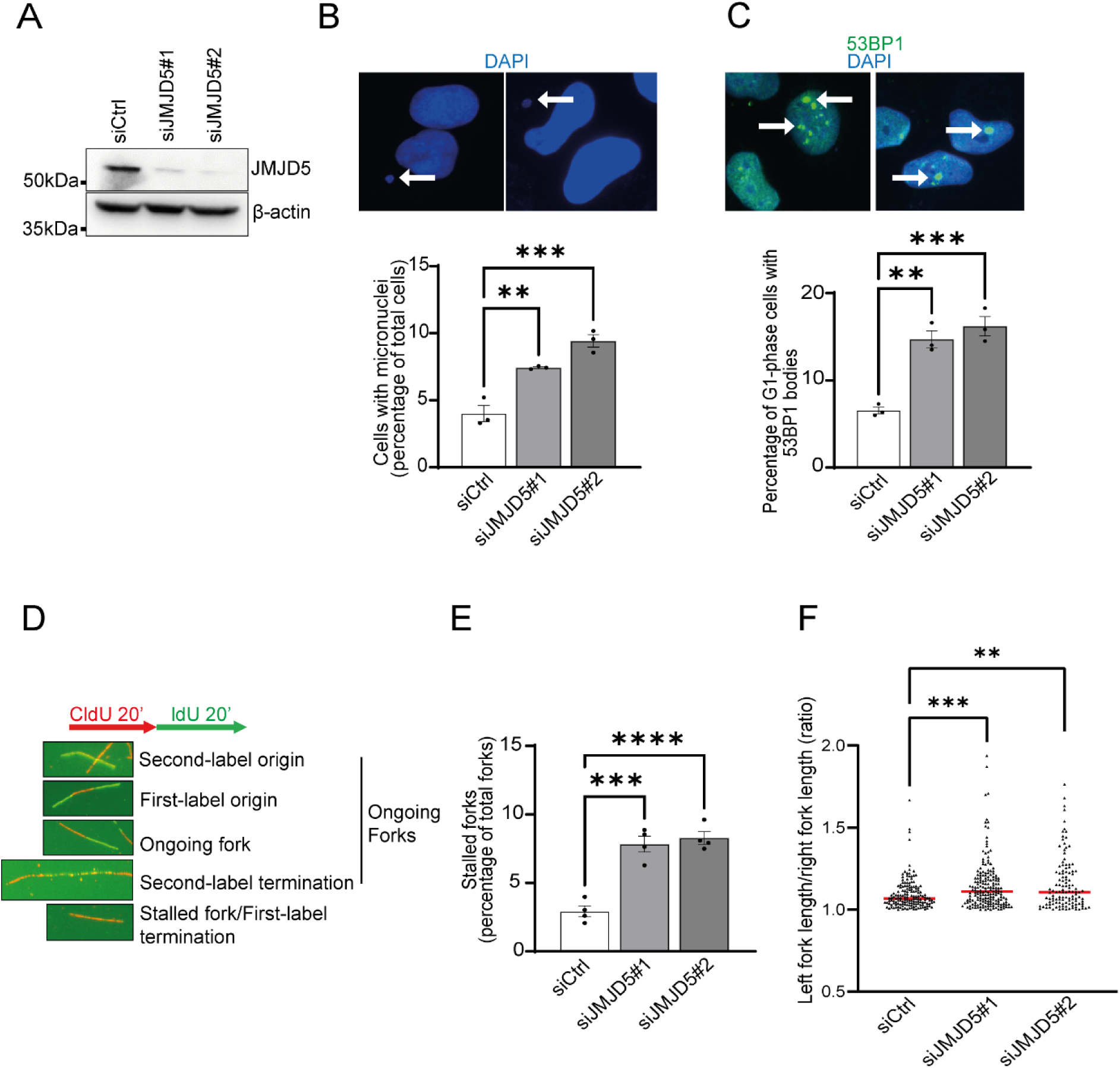
JMJD5 maintains replication fidelity in tumour cell lines. Lung adenocarcinoma A549 were treated with siRNA before quantification of RS markers including micronuclei, 53BP1 bodies, stalled forks and fork asymmetry. **(A)** A549 cells were transfected with the indicated siRNAs before western blot analysis to confirm JMJD5 knockdown. JMJD5 knockdown increased the number of **(B)** micronuclei and **(C)** 53BP1 bodies in G1 phase. **(D)** Single molecule DNA fibre analysis was performed by incubating cells with thymidine analogues CldU and IdU consecutively, before immunofluorescence detection (top). Examples of different fibre structures are shown (bottom). Both **(E)** stalled forks and **(F)** fork asymmetry was significantly increased following JMJD5 knockdown in A549 cells. Data represent mean ± SEM from 3 **(B, C,** and **F)** or 4 **(E)** independent experiments. For 53BP1 bodies, a minimum of 300 cells were counted per sample. For micronuclei, a minimum of 500 cells were counted per sample. For stalled forks a minimum of 250 forks were counted per condition per independent experiment. For fork asymmetry at least 50 structures were measured per sample per independent experiment. Statistical analyses used 1-way ANOVA with Bonferroni’s post hoc test (**B, C,** and **E**) or Kruskal-Wallis with Dunn’s correction (**F**); **P ≤ 0.01, ***P ≤ 0.001, ****P ≤ 0.0001.

Next, we used single molecule DNA fibre analysis to quantify stalled forks (Figure 1D), which increased during replication stress and following JMJD5 loss-of-function, as reported previously in primary dermal fibroblasts [13]. Stalled forks were elevated in A549 and U2OS cells in response to JMJD5 knockdown (Figure 1E and Supplemental Figure 2A), which was further exacerbated by the ribonucleotide reductase inhibitor hydroxyurea (HU) (Supplemental Figure 2B/C).

Unperturbed DNA replication proceeds bidirectionally from the same origin with comparable speeds, resulting in replication fork ‘symmetry’. Perturbed DNA replication may result in ‘fork asymmetry’, resulting from a slowing, pausing or collapsing of one of the two replication forks. Consistent with perturbed DNA replication, we observed significantly increased fork asymmetry following JMJD5 knockdown in A549 cells, both in the absence (Figure 1F) and presence (Supplemental Figure 2D) of HU. Likewise, we observed significant increases in fork asymmetry following JMJD5 knockdown in U2OS cells, both with and without HU treatment(Supplemental Figure 2E-F). Together, these results suggest that JMJD5 controls replication fidelity and genome stability in a variety of cell types and cancer models.

### JMJD5 is functionally inactivated by cancer variants

Tumour suppressor genes (TSGs) that control genome stability are functionally inactivated by a variety of mechanisms in cancer cells. Inherited variants in such genes can also cause susceptibility to cancer and early onset of the disease. Interestingly, two siblings affected by the previously described neurodevelopmental disorder inherited one of their bilateral inactivating JMJD5 mutations (C123Y) from their mother, who suffered from early onset breast cancer [13]. Her tumour genome did not contain known driver mutations (Illumina TruSight Oncology panel sequencing of 94 genes), raising the possibility that the JMJD5 missense variant contributed to her disease. Whether somatic JMJD5 tumour mutations also exist, and the potential role of these in RS, GIN, and tumourigenesis, is not known.

To investigate whether *JMJD5* is mutated in cancer, we mined tumour sequencing databases (Supplementary Table 1) and plotted variants against the JMJD5 protein sequence (Figure 2A). Missense substitutions, truncations and in-frame deletions were observed throughout the enzyme and in all three structural motifs including the N-terminal domain, JmjC extension and JmjC catalytic domain. Consistent with potential functional consequences, mutations were also recurrent and clustered in hotspots (Figure 2A). To investigate the impact of tumour-associated mutations on JMJD5 function, we cloned a panel of 28 cancer mutants, focussing on variants affecting conserved residues in key structural regions and with predicted functional impact (Supplementary Table 1, 2, Supplemental Figure 3). To investigate whether any of these variants compromised JMJD5 protein expression, we transfected HeLa cells with HA-tagged JMJD5 cancer mutants. Western blot analysis indicated that 12/28 variants negatively affected JMJD5 expression with six variants having a severe impact on protein expression. (Figure 2B, Supplementary Table 2). Next, we tested whether any of the variants negatively impacted JMJD5 catalytic activity using recombinant JMJD5 cancer mutants and an *in vitro* hydroxylation assay (as previously described [13]). Half of the cancer mutants displayed significantly reduced hydroxylase activity in comparison to WT JMJD5 (Figure 2C) (Supplementary Table 2). Overall, these data indicate that cancer mutants negatively impact JMJD5 expression and hydroxylase activity. Whether the apparently benign mutants (Supplementary Table 1) negatively affect function, but through alternative mechanisms, remains unclear.

**Figure 2.**
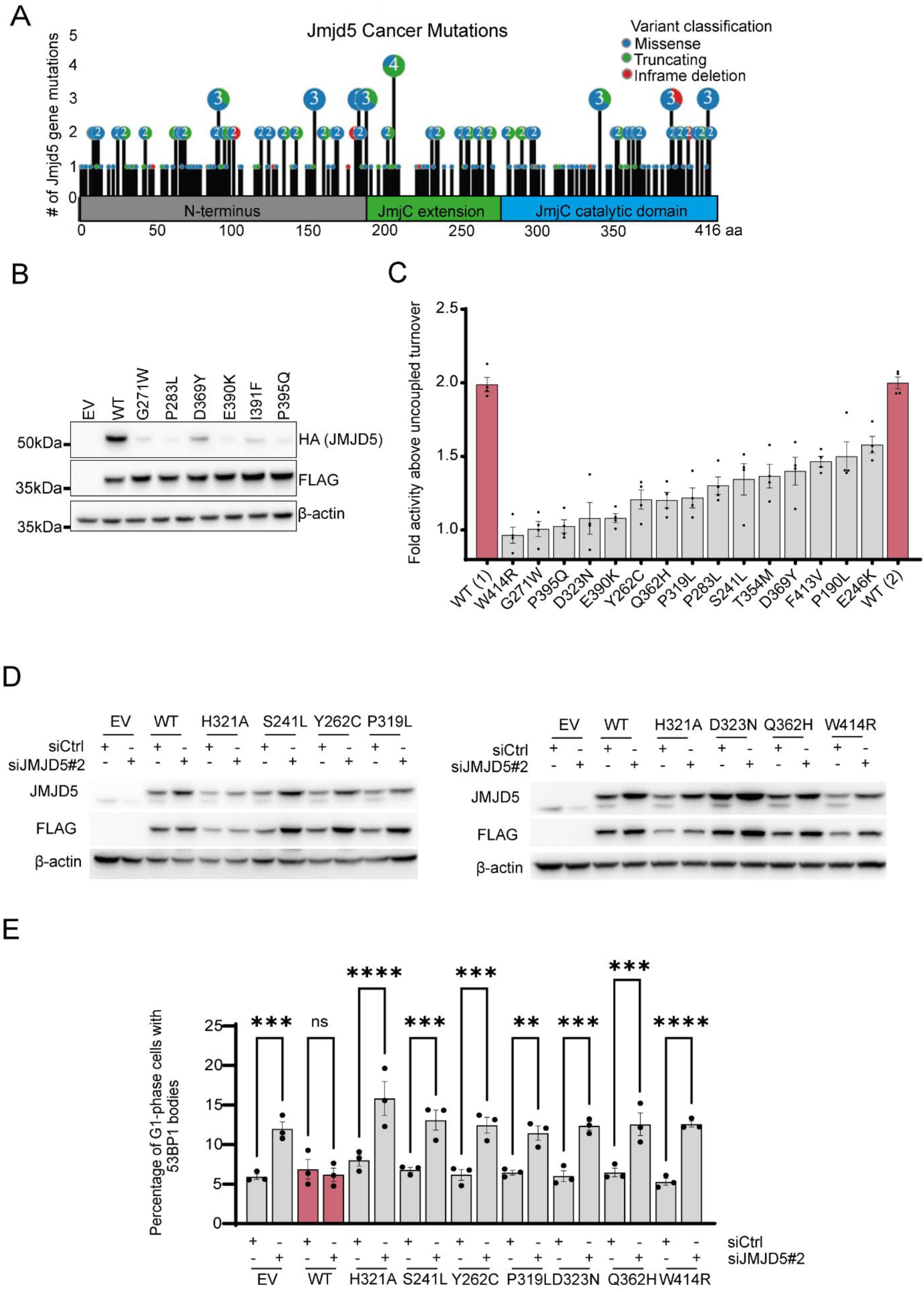
*JMJD5* cancer mutations reduce expression, catalytic activity and impair rescue of replication fidelity associated with JMJD5 knockdown. **(A)** Full list of *JMJD5* cancer mutations taken from online databases COSMIC and cBioPortal plotted against the primary sequence of JMJD5. **(B)** HeLa cells were transfected with the indicated HA-JMJD5 constructs for 48 h and JMJD5 expression examined by western blotting. FLAG-JMJD7 was used as a transfection control **(C)** GST-tagged WT JMJD5 and cancer variants were expressed in *E*. *coli* and purified before in vitro activity assays. Activity was monitored using the Succinate-Glo assay, which measures succinate production. Two separate WT preparations were included in each assay for comparison. Data represent mean ± SEM from 4 independent experiments. All variants shown had a statistically significant reduction in activity compared to WT JMJD5. **(D)** and **(E)** pTIPZ-FLAG JMJD5 A549 cell lines expressing the indicated JMJD5 cancer variants were treated with 10 ng/mL doxycycline for one week prior to transfection with siCtrl or siJMJD5#2. Cells were fixed after 72 h and 53BP1 bodies quantified in G1 phase cells. Re-expression of WT JMJD5 reduced levels of 53BP1 bodies in JMJD5 knockdown cells. In contrast, re-expression of the catalytically dead H321A mutant, or cancer variants shown to affect JMJD5 activity were unable to reduce 53BP1 bodies. Data represent mean ± SEM from 3 **(E)** or 4 **(B)** independent experiments. For 53BP1 bodies analysis a minimum of 250 G1 phase cells were counted per sample in each experiment. **(D)** western blot validation of JMJD5 reconstitution from panel **(E)**. Statistical analysis used 1-way ANOVA with Bonferroni’s post hoc test; ns not significant, **P ≤ 0.01, ***P ≤ 0.001, ****P ≤ 0.0001.

### JMJD5 cancer variants are unable to restore normal replication fidelity

Our previous work demonstrated that a disease-associated missense variant (C123Y) impaired JMJD5 hydroxylase activity, and that the role of JMJD5 in replication fidelity is activity-dependent [13]. Therefore, we hypothesized that cancer-associated missense variants that impair hydroxylase activity would also impact the role of JMJD5 in replication fidelity. To test this, we focussed on six JMJD5 cancer variants that were expressed normally but had reduced activity (S241L, Y262C, P319L, D323N, Q362H and W414R) (Figure 2B-C). These mutants were cloned into a doxycycline-inducible lentiviral vector for conditional re-expression following JMJD5 knockdown (because our siRNA sequences target the JMJD5 mRNA 3’UTR, the reconstituted cDNAs are resistant to knockdown). First, we confirmed successful knockdown of endogenous JMJD5 and reconstitution of exogenous wildtype JMJD5, an inactive mutant (H321A), or the JMJD5 cancer variants (Figure 2D). Next, we tested the ability of the cancer variants to prevent replication stress induced by endogenous JMJD5 knockdown. As hypothesized, cancer variants that reduced JMJD5 hydroxylase activity were also unable to restore normal levels of 53BP1 bodies (Figure 2E) or micronuclei (Supplemental Figure 4).

### The role of JMJD5 in replication fidelity is epistatic with its interactor, RCCD1

How JMJD5 inactivation causes replication stress in tumour cells, and whether it acts in isolation, or as part of a wider complex or pathway, is not yet clear. To begin exploring this, we initially considered the involvement of known JMJD5 interacting proteins. Previous studies have shown that JMJD5 interacts with ‘RCC1 domain-containing protein 1’ (RCCD1) [20, 21], a poorly characterised seven-bladed β-propeller protein. Like JMJD5, RCCD1 has also been implicated in a variety of cancer types [22–25]. To investigate whether RCCD1 might act in the same cellular pathway as JMJD5, we first tested whether RCCD1 depletion also caused replication stress. RCCD1 knockdown in A549 (Figure 3A-C) and U2OS (Supplemental Figure 5) caused an increase in 53BP1 bodies and micronuclei. Furthermore, single molecule DNA fibre analysis demonstrated that RCCD1 depletion increased fork stalling and asymmetry in both the absence (Figure 3D-E) and presence (Supplemental Figure 6) of HU. Similar results were obtained in U2OS cells (Supplemental Figure 7).

**Figure 3.**
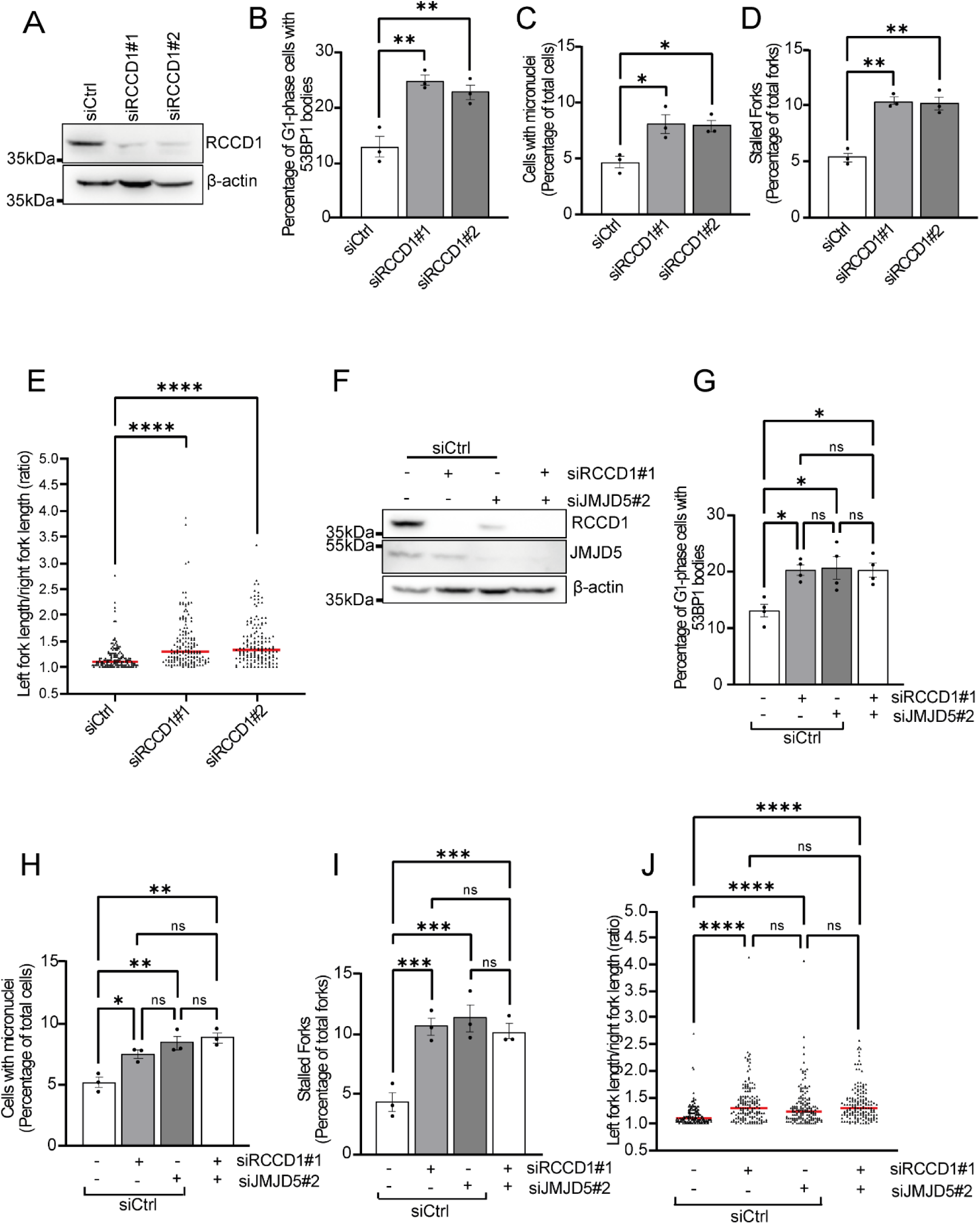
RCCD1 depletion increases markers of replication stress epistatic with JMJD5. A549 cells were treated with siRNA and multiple markers of replication stress were quantified including micronuclei, 53BP1 bodies, stalled forks and fork asymmetry. **(A)** A549 cells were transfected with indicated siRNAs and western blotted using the indicated antibodies to confirm RCCD1 knockdown. siRNA knockdown of RCCD1 using two independent sequences increased the number of **(B)** 53BP1 bodies, **(C)** micronuclei, **(D)** stalled forks and **(E)** asymmetric forks. **(F)** Successful knockdown of JMJD5 and RCCD1 using indicated siRNAs in A549 cells was confirmed via western blot. Cells were harvested 72 h post transfection and analysed using the indicated antibodies. The JMJD5 and RCCD1 increased replication stress phenotypes are epistatic as indicated by no additional increase in **(G)** 53BP1 bodies, **(H)** micronuclei, **(I)** stalled forks and **(J)** fork asymmetry following concurrent knockdown. Data represent mean ± SEM from 3 **(B, C, D, E, H, I** and **J)** or 4 **(G)** independent experiments. For 53BP1 bodies, a minimum of 300 cells were counted per sample. For micronuclei, a minimum of 500 cells were counted per sample. For stalled forks a minimum of 250 forks were counted per condition per independent experiment. For fork asymmetry at least 50 structures were measured per sample per independent experiment. Statistical analyses used 1-way ANOVA with Bonferroni’s post hoc test (**B, C, D, G, H** and **I**) or Kruskal-Wallis with Dunn’s correction (**E** and **J**); *P ≤ 0.05, **P ≤ 0.01, ***P ≤ 0.001, ****P ≤ 0.0001.

To test if JMJD5 and RCCD1 act in the same pathway controlling replication fidelity, we first aimed to explore their epistatic relationship to one another following single or combined siRNA knockdown (Figure 3F). We observed that simultaneous knockdown of JMJD5 and RCCD1 in A549 cells did not increase levels of 53BP1 bodies, micronuclei, stalled forks, or fork asymmetry, beyond that observed with single knockdown (Figure 3G-J). Similar results were obtained for 53BP1 bodies and micronuclei in U2OS cells (Supplemental Figure 8). Overall, these observations indicate that JMJD5 and RCCD1 likely act in the same pathway to maintain replication fidelity.

Whilst validating knockdowns in the epistasis experiments described above, we also noticed a potential inter-dependency in JMJD5 and RCCD1 protein expression. In A549 cells we observed that JMJD5 siRNA knockdown resulted in reduced RCCD1 expression (Figure 3F), whilst in U2OS cells we noticed a reduction in JMJD5 expression following RCCD1 siRNA knockdown (Supplemental Figure 8). These observations, coupled with the apparent epistatic relationship outlined above, led us hypothesise that JMJD5 and RCCD1 are obligate binding partners that need to act in a complex together for optimal expression and function in replication fidelity. To explore this further, we first characterised the JMJD5:RCCD1 interaction in more detail.

### JMJD5 and RCCD1 form a stoichiometric 1:1 heterodimer dependent on key electrostatic interactions

We began by overexpressing JMJD5 in the presence or absence of overexpressed RCCD1, observing that co-expression of RCCD1 increased the levels of JMJD5 (Figure 4A). Furthermore, Coomassie staining of SDS-PAGE gels following co-immunoprecipitation demonstrated that JMJD5 and RCCD1 form a stable complex (Figure 4B) and a partially purified complex eluted as a single peak with an apparent molecular weight of ∼90kDa consistent with the formation of a 1:1 heterodimer between JMJD5 (∼47kDa) and RCCD1 (∼40kDa) (Supplemental Figure 9)

**Figure 4.**
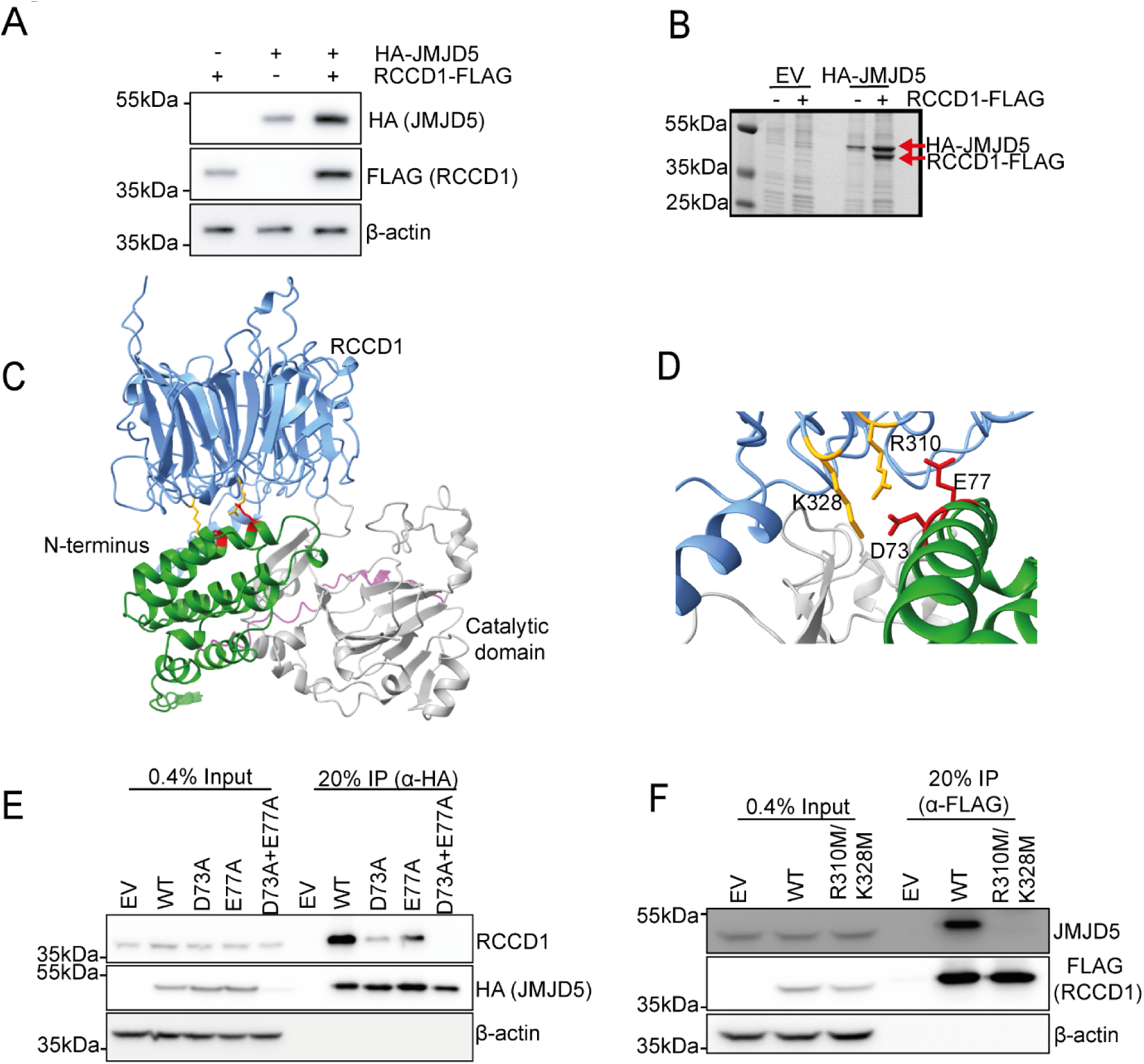
JMJD5 and RCCD1 form a 1:1 heterodimer facilitated through D73A/E77A JMJD5. **(A)** Expression of JMJD5 and RCCD1 is optimal when co-transfected. HEK293T cells were transfected with either pIHZ RCCD1-FLAG, pEF6 HA-JMJD5, or both, before western blotting with the indicated antibodies. **(B)** JMJD5 and RCCD1 form a stoichiometric 1:1 complex. HEK293T cells were transfected with indicated pEF6 constructs (EV or HA-JMJD5) and pIHZ RCCD1-FLAG followed by anti-HA immunoprecipitation (IP). The immunoprecipitated samples were then separated using SDS-PAGE and stained using Coomassie. **(C)** AlphaFold3 prediction showed RCCD1 (blue) interacts with the N-terminus (green) of JMJD5 which is connected to the catalytic domain (grey) through a flexible linker (pink) **(D)** AlphaFold Multimer prediction supports the identification of JMJD5 D73/E77 (red) and RCCD1 R310/K328 (orange) forming the binding interface between the two proteins. **(E)** RCCD1 is unable to bind to JMJD5 D73A/E77A. Lysates from HEK293T cells transfected with control pEF6 (EV), pEF6-HA-JMJD5(WT) or the indicated experimental alanine mutations pEF6-HA-JMJD5(D73A, E77A or D73A+E77A) were subject to anti-HA immunoprecipitation and western blotting with the indicated antibodies. **(F)** Experimental mutation of R310 and K328 to methionine does reduce binding to JMJD5. Lysate from HEK293T cells transfected with control pIHZ (EV), pIHZ-RCCD1-FLAG(WT) and pIHZ-RCCD1-FLAG(R310M/K328M) were anti-FLAG immunoprecipitated (IP) followed by western blotting with the indicated antibodies.

Available empirical structural information for these proteins is currently limited to X-ray structures of the Jmjd5 catalytic domain[12]. To identify amino acids that mediate JMJD5/RCCD1 binding we used AlphaFold to generate a model of the complex (Figure 4C). Within this predicted structure, the catalytic domain (CD) closely resembles that of extant crystal structures with a core double stranded β-helix (DSBH) that forms the active site cleft, elaborated by the classical N-terminal extension (NTE) that wraps around and extends the DSBH sheet. Consistent with our earlier modelling studies [13], the JMJD5 N-terminal domain (NTD) forms a helical stack structure reminiscent of tetratricopeptide repeats (TPR). Its two C-terminal helices (α4/5) pack exclusively against the CD N-terminal extension through contributing to a wedge-shaped arrangement for the full-length enzyme. As expected from its sequence homology with RCC1 [26], RCCD1 adopts a 7-bladed β-propellor architecture and interacts with α2 of the JMJD5 NTD through a surface centred on an unusual cluster of exposed Trp residues (Figure 4C) through a largely hydrophobic interface further stabilised by electrostatic/salt-bridging interactions between D73 and E77 from JMJD5 and R310 and K328 on RCCD1 (Figure 4D).

Supporting the veracity of the AlphaFold model, substitution of either D73 or E77 significantly reduced the interaction of JMJD5 with RCCD1 in co-IP assays, whilst a D73A/E77A double mutant completely ablated binding (Figure 4E). To test the impact of mutating the corresponding residues in RCCD1, we used methionine substitutions to maintain hydrophobic packing of the aliphatic portions of the R310 and K328 side-chains against surface-exposed Trp residues, W220 and W326 respectively. Consistent with the AlphaFold predictions, an R310M/K328M mutant completely blocked the interaction of RCCD1 with JMJD5 (Figure 4F). Overall, these and previous observations validate the structural prediction and confirm the critical contribution of electrostatic interactions to JMJD5:RCCD1 complex stability.

Disruption of critical interactions in tumour suppressor complexes can be positively selected during cancer development. If the interaction of JMJD5 and RCCD1 is essential for maintaining replication fidelity and suppressing tumourigenesis, we hypothesised that some of the interacting residues identified above would be mutated in tumour samples and that these would disrupt the JMJD5:RCCD1 interaction. Interrogating the cBioPortal and COSMIC databases identified a JMJD5 E77K mutation in a malignant melanoma sample that would be predicted to disrupt one of the critical electrostatic interactions with RCCD1. Other cancer mutants were also identified in close proximity to D73 and E77 (V70L, S75F and L79P) (Supplementary table 1). Whereas V70L and S75F substitutions did not impact the JMJD5:RCCD1 interaction under the conditions tested, the L79P and E77K variants did (Supplemental Figure 10). Similarly, we identified RCCD1 missense substitutions in residues that are critical for the JMJD5 interaction, including R310W/Q (Ewings Sarcoma and Skin Squamous Cell Carcinoma) and K328N (uterine carcinoma) (Supplementary Table 3). The R310W and K328N mutations significantly compromised the ability of JMJD5 to interact with RCCD1 (Supplemental Figure 10).

These data suggest that critical determinants of the JMJD5:RCCD1 interaction predicted from the AlphaFold model may be altered in human tumours, thus highlighting the importance of understanding the function of the JMJD5:RCCD1 in greater detail.

### JMJD5:RCCD1 complex formation is essential to maintain replication fidelity in tumour cell lines

Having identified experimental and disease mutants that disrupt the JMJD5:RCCD1 interaction, we next sought to leverage these to test the importance of complex formation for maintaining replication fidelity in tumour cells, initially focussing on the JMJD5 D73A/E77A mutant. To do so, we moved to the ‘rescue’ system described above. Western blot analyses confirmed successful knockdown and re-expression of wildtype and H321A JMJD5, although comparable re-expression of the D73A/E77A mutant required a 100-fold higher doxycycline concentration (consistent with the importance of the RCCD1 interaction for JMJD5 expression) (Figure 5A). Whereas reconstitution with wildtype JMJD5 was able to fully restore control levels of 53BP1 bodies (Figure 5B), the D73A/E77A RCCD1-binding mutant was not, behaving more similarly to the functionally inactive H321A mutant. Quantification of other replication stress markers confirmed that reconstitution with D73A/E77A JMJD5 was unable to restore control levels of micronuclei (Supplemental Figure 11), stalled forks (Figure 5C) or fork asymmetry (Figure 5D).

**Figure 5.**
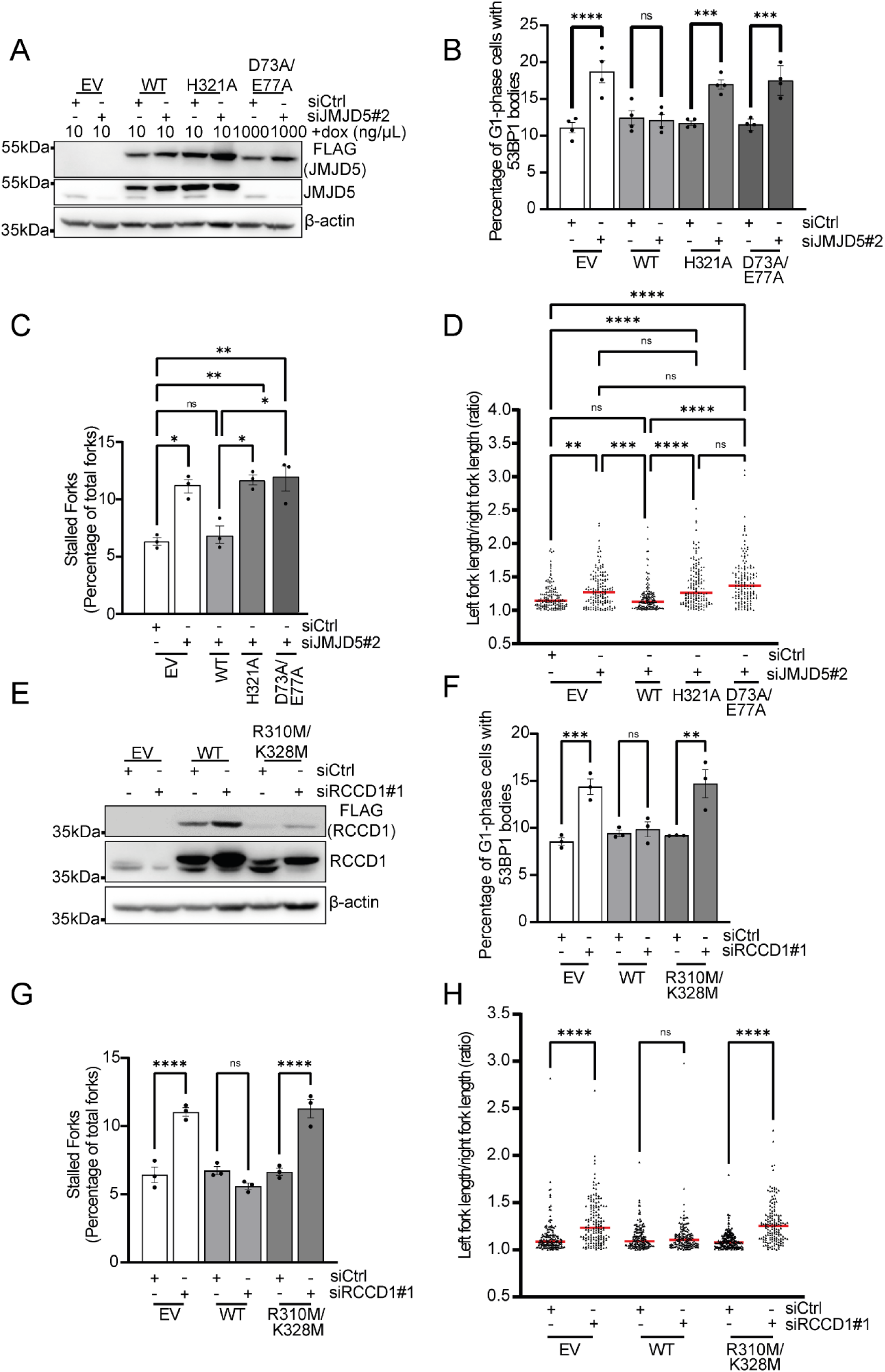
JMJD5 and RCCD1 binding mutants are unable to rescue siRNA induced increases in markers of replication stress. pTIPZ-JMJD5 or pIHZ-RCCD1 A549 cells were transfected with indicated siRNAs before quantification of 53BP1 bodies, stalled forks and fork asymmetry. **(A)** The indicated pTIPZ-JMJD5 A549 cell lines (EV, WT, H321A and D73A/E77A) were treated with indicated concentrations of doxycycline for one week prior to transfection with siCtrl or siJMJD5#2. Cells were harvested after 72h and JMJD5 knockdown was confirmed via western Blot with the labelled antibodies. Note that expression of D73A/E77A was not detected using the JMJD5 antibody as these mutations presumably destroy the antigen. pTIPZ-JMJD5 WT but not H321A or D73A/E77A was able to rescue increases in **(B)** 53BP1 bodies, **(C)** stalled forks and **(D)** fork asymmetry caused by siJMJD5. **(E)** pIHZ-RCCD1 (EV, WT, R310M/K328M) were transfected with the indicated siRNAs and knockdown of RCCD1 was confirmed via western Blot. pIHZ-RCCD1 WT but not the JMJD5 binding mutant R310M/K328M showed rescue of siRCCD1#1 induced increases in **(F)** 53BP1 bodies, **(G)** stalled forks and **(H)** fork asymmetry. Data represent mean ± SEM from 3 **(C, D, F, G and H)** or 4 **(B)** independent experiments. For 53BP1 bodies, a minimum of 300 cells were counted per sample. For stalled forks a minimum of 250 forks were counted per condition per independent experiment. For fork asymmetry at least 50 structures were measured per sample per independent experiment. Statistical analyses used 1-way ANOVA with Bonferroni’s post hoc test (**B, C, F** and **G**) or Kruskal-Wallis with Dunn’s correction (**D** and **H**); *P ≤ 0.05, **P ≤ 0.01, ***P ≤ 0.001, ****P ≤ 0.0001.

To test if the reciprocal was also true, we generated A549 cells constitutively expressing wildtype RCCD1 or the JMJD5 binding mutant R310M/K328M (Figure 5E). Similarly to the D73A/E77A JMJD5 mutant reconstitution experiments above, the R310M/K328M mutant was less well expressed than wildtype RCCD1, consistent perhaps with the importance of the interaction with JMJD5 for optimal protein expression. Despite the reduced expression of the R310M/K328M mutant from this constitutive vector, we noted it was still expressed to similar or slightly higher levels than endogenous RCCD1 (Figure 5E). Therefore, we reasoned it could still be used to restore physiological levels of RCCD1 expression that could be used to test the functional competence of this mutant. Whereas wildtype RCCD1 was able to restore replication stress markers to control levels, the JMJD5 binding mutant was not (Figure 5F-H and Supplemental Figure 12).

Although our data demonstrates a clear role for JMJD5:RCCD1 complex formation in maintaining replication fidelity in tumour cells, the precise nature of its role requires further investigation.

### The JMJD5:RCCD1 complex interacts with HR factors and is required for restart of stalled replication forks

To explore the role of JMJD5 and RCCD1 in replication fidelity and human disease in more detail, we were motivated to identify their most proximal biology using mass spectroscopy (MS)-based interactomics. Early attempts using single JMJD5 or RCCD1 pulldowns did not yield high fidelity assignment of nuclear proteins known to play a role in replication or associated biology. In light of this, and our findings demonstrating the critical importance of the JMJD5:RCCD1 interaction for their function, we reasoned that new proteomic screens were warranted using the JMJD5:RCCD1 complex as bait. To this end we used a multicistronic doxycycline-inducible vector containing HA-JMJD5 and RCCD1-FLAG cDNAs separated by a P2A sequence to generate two separately expressed proteins that we confirmed formed a 1:1 complex (Supplemental Figure 13).

Next, we considered the need to enrich for relevant JMJD5:RCCD1 interactors by treating cells with a relevant stressor. We opted for a prolonged HU treatment because it induces replication stress and DNA damage that activates homologous recombination (HR) [27]. The rationale for this was that, in addition to its role in replication fidelity, JMJD5 has also been implicated in the late stages of HR [28], similar to several other factors implicated in both the RS and DNA damage responses [29]. MS/MS sequencing of large-scale anti-FLAG immunoprecipitations under these conditions identified more than fifty proteins that were specific to JMJD5:RCCD1 pulldowns and that met our filtering criteria (Supplementary Table 4). To triage the hits, we screened the candidates for HR and/or replication factors and identified two RAD51 paralogues known as RAD51C and XRCC3. RAD51C and XRCC3 form a complex known as CX3 which has been implicated in late-stage HR [30] and replication fork restart [31]. Independent co-immunoprecipitation experiments in HEK293T cells following 24 hours of HU treatment confirmed that JMJD5:RCCD1 interacts specifically with RAD51C and XRCC3, but not RAD51, other RAD51 paralogues (RAD51B), or RPA (Figure 6A). Purified JMJD5:RCCD1 complex also interacted specifically with recombinant CX3 complex *in vitro* (Supplemental Figure 14).

**Figure 6.**
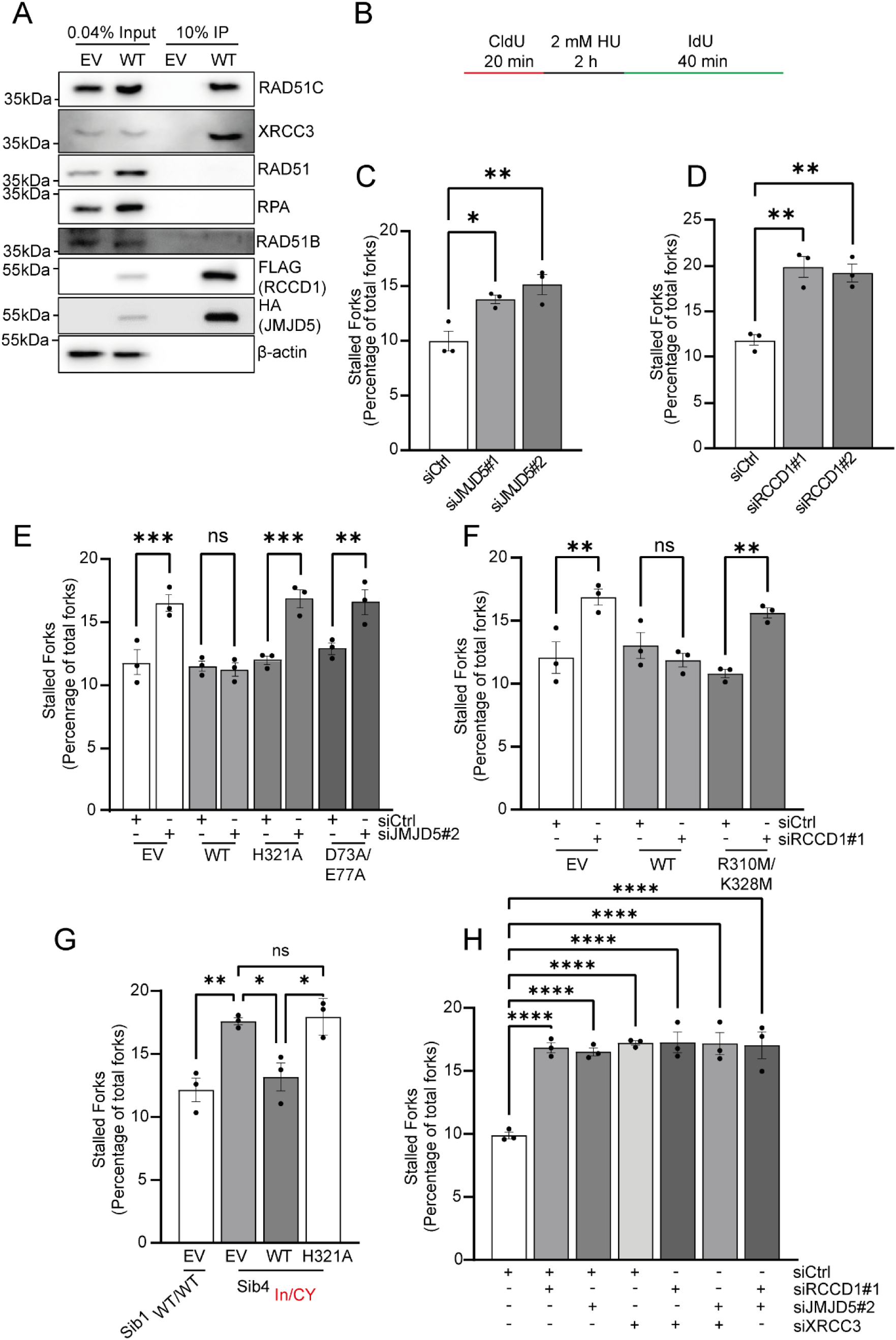
The JMJD5:RCCD1 heterodimer interacts with the CX3 complex of RAD51 paralogs to facilitate normal restart of stalled replication forks. **(A)** The JMJD5:RCCD1 heterodimer interacts with the CX3 complex of RAD51 paralogs. HEK293T cells expressing doxycycline inducible pTIPZ EV or WT HA-JMJD5-P2A-RCCD1-3XFLAG were treated with doxycycline for 48 h before 2 mM HU treatment for 24 h. Resulting protein lysates were anti-FLAG immunoprecipitated (IP) followed by western blotting with the indicated antibodies. **(B)** Modified DNA fibre assay. To measure fork restart CldU/IdU was pulse-labelled for indicated times with HU treatment (2 mM, 2 h). siRNA knockdown of both **(C)** JMJD5 and **(D)** RCCD1 in A549 cells significantly increases the number of stalled forks associated with deficient replication fork restart. Formation of the JMJD5:RCCD1 heterodimer is required for restart of stalled replication forks with **(E)** Reconstitution of WT JMJD5 but not catalytically dead (H321A) or RCCD1 binding mutant (D73A/E77A) and **(F)** reconstitution of WT RCCD1 but not the JMJD5 binding mutant (R310M/K328M) able to rescue observed increases in HU induced deficient replication fork restart in A549 cells. **(G)** Reconstitution of active JMJD5 only in Sib4_In/CY_ could rescue the observed increase in deficient replication fork restart. **(H)** Observed JMJD5/RCCD1 knockdown associated replication fork restart defects are epistatic with functional inactivation of the CX3 complex of RAD51 paralogs in A549 cells. For stalled forks associated with deficient replication fork restart a minimum of at least 50 structures were counted per condition per independent experiment. Data represents mean ± SEM from 3 **(C, D, E, F, G, and H)** independent experiments. Statistical analyses used 1-way ANOVA with Bonferroni’s post hoc test **(C, D, E, F, G, and H).** *P ≤ 0.05, **P ≤ 0.01, ***P ≤ 0.001, ****P ≤ 0.0001. ns = non-significant.

The CX3 complex has an important role in replication fork restart [31]. This raised the possibility that the function of the JMJD5:RCCD1 complex in replication fidelity and disease might be related to restarting stalled replication forks. To investigate this, we used a modified version of the DNA fibre assay (Figure 6B) [31]. Consistent with our hypothesis, siRNA knockdown of either JMJD5 (Figure 6C) or RCCD1 (Figure 6D) caused a significant increase in stalled forks, which in this assay indicates a deficit in replication fork restart. We next engaged our JMJD5 and RCCD1 reconstitution models, noting that both formation of the JMJD5:RCCD1 complex (Figure 6E and 6F) and the catalytic activity of JMJD5 (Figure 6E) are essential for replication fork restart.

Our earlier study reporting the role of JMJD5 in replication fidelity followed the identification of JMJD5 loss-of-function variants as the causal basis of a novel neurodevelopmental disorder [13]. In light of our findings in tumour cell lines, we considered the possibility that replication fork restart was also defective in our neurodevelopmental disorder patients. Primary cells derived from an affected patient carrying biallelic inactivating JMJD5 variants displayed a deficit in replication fork restart (Supplemental Figure 15), which could be rescued by reconstitution of catalytically active JMJD5 (Figure 6G).

Finally, we tested the hypothesis that JMJD5, RCCD1, and CX3 function in the same pathway in replication fork restart using epistasis analysis. Knockdown of XRCC3 was used to selectively inhibit the CX3 complex, as reported previously [31] (Supplemental Figure 16). Interestingly, co-depletion of either RCCD1/XRCC3 or JMJD5/XRCC3 did not exacerbate the replication fork restart deficit beyond that observed following knockdown of each factor in isolation (Figure 6H). This suggests that both heterodimeric complexes, JMJD5:RCCD1 and RAD51C:XRCC3, act in the same pathway controlling replication fork restart, and that this pathway underlies our observations of a role of the JMJD5:RCCD1 complexes in replication fidelity, neurodevelopmental disorders, and cancer.

## Discussion

The findings presented here build on our prior work (REF patient paper) to further evidence the importance of the JMJD5 hydroxylase in human health and disease. We show that JMJD5 variants identified in human tumours impair protein expression (Figure 2B), and enzymatic activity (Figure 2C), and demonstrate that JMJD5 function is required to maintain replication fidelity in tumour cell lines (Figure 2E). We additionally demonstrate that JMJD5 forms a heterodimer with a cancer-associated seven-bladed β-propeller protein called RCCD1 (Figure 4), which we show also maintains replication fidelity. Critically, the role of RCCD1 in replication is dependent on its interaction with JMJD5 (Figure 3 and 5). Proteomic pulldowns using the JMJD5:RCCD1 complex identified a novel interaction with the RAD51 paralogue CX3 complex in tumour cell lines (Figure 6A), which led to us uncover an epistatic relationship between these two enigmatic complexes in replication fork restart (Figure 6H). Together with our observation of defective replication fork restart in JMJD5 patient cells (Figure 6G and Supplemental Figure 15), our work has the potential to unify the role of the JMJD5:RCCD1 complex in neurodevelopmental disorders and cancer.

The disruption of the JMJD5:RCCD1 interaction by damaging cancer-associated mutations (Supplemental Figure 10) suggests a previously unrecognised tumour suppressor axis involving a protein hydroxylase which, as an enzyme class, has not previously been widely studied in the context of cancer. Although the frequency of JMJD5 and RCCD1 *missense mutations* across all cancers appears relatively low, the full extent of JMJD5:RCCD1 functional inactivation in tumours is not yet known. Future work will address this by investigating alternative mechanisms of JMJD5:RCCD1 inactivation including, for example, copy number variation, epigenetic silencing, single nucleotide polymorphisms (SNPs) in transcription regulatory sequences, aberrant mRNA splicing, and functional inactivation of the proteins themselves.

Through a series of genome-wide and transcriptome-wide association studies, RCCD1 has been independently proposed as a novel TSG. A genome wide association study identified a novel SNP (rs2290203) located in an intron of a gene 5,712 bp downstream of *RCCD1* [32], which correlated with reduced RCCD1 mRNA expression and increased breast cancer risk. Additional RCCD1-associated SNPs at the *15q26.1* genome location have been reported in ovarian cancer (rs8037137) [25], breast cancer (rs3826033, rs2290202 and rs17821347) [23] and pancreatic cancer (rs8028409) [33], all of which correlate with reduced mRNA expression and increased disease risk.

Similarly, [34] identified risk variants in which increased RCCD1 expression correlated with a reduced risk of breast cancer. Whether the reduced RCCD1 expression in these contexts causes reduced JMJD5 protein expression (as reported here, Figure 4) and function, and the potential impact of this on replication fork restart and genome stability, is not yet clear.

The tumour microenvironment has the potential to also impair JMJD5:RCCD1 function, given the unique cofactor and nutrient dependencies of 2OG oxygenases [35]. Oncometabolites such as fumarate, succinate, and D-2-hydroxyglutarate (D-2HG) produced by mutations in FH, SDH, and IDH1/2, respectively, can competitively inhibit 2OG oxygenases due to their structural similarity to 2OG [5]. This raises the possibility that JMJD5 activity may be similarly compromised in metabolically reprogrammed tumours.

Hypoxia represents another potential mechanism of JMJD5 inactivation. While HIF prolyl hydroxylases are well-characterised oxygen sensors, the oxygen sensitivity of JmjC-only hydroxylases such as JMJD5 remains poorly defined [5]. Of interest to the work presented here, and the potential oxygen-sensitivity of JMJD5, hypoxic stress is known to suppress homologous recombination, impair DNA replication, and promote genomic instability [36]. Whether JMJD5 inhibition contributes to hypoxia-induced replication defects remain to be determined, but our findings suggest this is a promising avenue for future investigation with broad clinical relevance.

Whilst the evidence presented here demonstrates a clear role for the JMJD5:RCCD1 heterodimer in replication fidelity (Figure 4 and 5), further work is required to uncover the precise role of each protein. For example, the presence of RCCD1 raises questions about its potential role in regulating JMJD5 catalytic activity, substrate binding, and subcellular localisation. To lay the groundwork for detailed future investigation of these possibilities, we identified the interactome of the JMJD5:RCCD1 complex, for the first time, by developing and applying a P2A-based dual expression system (Figure 6). The novel interaction we describe between the JMJD5:RCCD1 and CX3 heterodimers (Figure 6) may be consistent with a role in a specific aspect of replication fork biology. CX3 is one of two separate RAD51 paralog complexes that also include ‘BCDX2’. Whereas CX3 plays a role in replication fork restart, as described above, BCDX2 is required for replication fork reversal [31]. The absence of RAD51B in JMJD5:RCCD1 pulldowns (Figure 6) suggests the interaction with RAD51 paralogs is specific to CX3, pointing to a role for JMJD5:RCCD1 in replication fork restart over reversal. Consistent with this, we did not observe a decrease in replication fork speed in JMJD5 mutant patient cell lines [13], which would ordinarily be indicative of a defect in replication fork reversal [37]. Rather we observed that JMJD5, RCCD1 and, importantly, the interaction between them, is required for replication fork restart consistent with a potential functional relationship with the CX3 complex. In line with this, we demonstrate that the JMJD5:RCCD1 and CX3 heterodimers are epistatic to one another with respect to defective replication fork restart (Figure 6). Beyond replication fork restart, the CX3 complex also plays an important role in DNA damage repair through the Homologous Recombination (HR) pathway [30, 38]. Therefore, the interaction with CX3 may also suggest a potential role for the JMJD5:RCCD1 complex in HR. Interestingly, JMJD5 loss of function has been shown to hypersensitise to ionising radiation and delay the resolution of RAD51 foci in C. elegans [28, 30], which aligns with similar phenotypes observed following CX3 depletion in other models [30]. Future work will focus on understanding the biochemical relationship between CX3 and JMJD5:RCCD1 in more detail, including whether this relates to the hydroxylase activity of JMJD5, and the molecular mechanism by which this regulates replication fork restart and HR.

The identification here of a novel cancer-associated pathway raises the possibility of developing new therapeutic approaches. We recently developed novel JMJD5 inhibitors and demonstrated that they cause increased replication stress in tumour cell lines [39], consistent with the importance of JMJD5 catalytic activity for replication fidelity and genome stability (Figure 2 and [13]). Inhibition of factors involved in these and related cellular processes often sensitises cells to genotoxic agents. Indeed, RAD51C and XRCC3 loss-of-function can sensitise tumour cells to a variety of genotoxic agents including PARP inhibitors and Cisplatin [40, 41]. It will, therefore, be of interest to determine whether inactivation of the JMJD5:RCCD1 complex, be that genetic, chemical, or biochemical, also sensitises tumour cells to genotoxic agents, and whether combination therapies might provide an additional new therapeutic opportunity.

In line with our evidence for damaging JMJD5:RCCD1 mutations in tumours, reports of reduced RCCD1 expression being linked to cancer predisposition, and the functional and physical interaction of JMJD5:RCCD1 with the CX3 complex, both RAD51C and XRCC3 have been implicated in cancer susceptibility and tumourigenesis [42–45]. Additionally, defects in the RAD51C component of the CX3 complex are the basis of the developmental disorder Fanconi Anaemia [42]. Whilst our previously reported clinical and cellular phenotypes for patients affected by damaging JMJD5 variants are not directly analogous to Fanconi Anaemia, they are both congenital disorders underpinned by increased genome instability [13, 42]. Our combined biochemical, clinical and functional data would predict that inherited variants that impair RCCD1 function (or other as yet unknown components of the pathway) may be the cause of some undiagnosed neurodevelopmental disorders.

In summary, we have provided further evidence of the disease relevance of the protein hydroxylase JMJD5, identifying other key members of a potential new tumour suppressor pathway involving RCCD1, RAD51C and XRCC3. In doing so, we have significantly expanded our knowledge of JMJD5 biology, and that of protein hydroxylases more broadly. Our work also highlights the potential importance of ‘obligate’ binding partners for the function of JmjC-hydroxylases, in this case RCCD1 and its role in the restart of stalled replication fork. The work supports the increasing evidence that JmjC-only 2-OG oxygenases are key modulators of human disease progression that warrant further investigation in both the discovery and translational research spaces.

## Materials and Methods

### DNA expression vectors

N-terminally HA epitope-tagged JMJD5 in pEF6 was generated as previously described [13]. N-terminally FLAG epitope-tagged doxycycline-inducible JMJD5 expression vectors, were created by PCR subcloning into modified lentiviral pTRIPZ (Dharmacon) vectors, in which the RFP cDNA was replaced with a modified multiple cloning site (termed pTIPZ, a gift from Matthew Cockman, Francis Crick Institute, London, UK). N-terminally glutathione S-transferase-tagged (GST-tagged) pGEX-4T-1 JMJD5 was generated as previously described [13]. C-terminally HA epitope-tagged RCCD1 expression vector was generate by cloning human *RCCD1* open reading frame (ORF) by PCR using Phusion High Fidelity DNA Polymerase (New England Biolabs) into pcDNA3.1 (Thermo Fisher). Lentiviral C-terminally FLAG-epitope tagged RCCD1 expression vector was generated by PCR subcloning into modified pGIPZ in which the GFP cDNA was replaced with a limited multiple cloning site and the puromycin resistance cDNA was replaced with hygromycin resistant cDNA (termed pIHZ; gift from Tencho Tenev, Institute of Cancer Research, London, UK). Both pEF6 HA-JMJD5 and pcDNA3.1 RCCD1-HA were used as templates for site-directed mutagenesis PCR reactions as previously described [13]. Following SDM PCR, successful RCCD1 clones were subcloned into pIHZ as above. The DNA sequence for HA-JMJD5-P2A-RCCD1-3XFLAG was commercially ordered from TransOmic and cloned into pTIPZ via restriction cloning.

### Cell treatments

siRNA transfections were performed using Lipofectamine RNAiMAX (Invitrogen) according to manufacturer’s instructions. All siRNAs were used at a working concentration of 30 nM. siRNA oligonucleotides were purchased from Sigma-Aldrich (Supplemental Figure 17). Control transfections were carried out using a negative control siRNA (Sigma-Aldrich SIC001), as previously described [13]. DNA plasmid transfection was performed using FuGENE (Promega) according to the manufacturer’s instructions. N-terminally FLAG epitope-tagged JMJD5 expression was induced using doxycycline, 24 hours after plating of stable cell lines. Concentration and duration of doxycycline treatment is indicated for each experiment. 1 mM hydroxyurea treatment was used where indicated.

### Cell culture

A549 (CCL-185), HeLa (CLL-2) and HEK293T (CRL-3216) cell lines were purchased from ATCC. U2OS cells were a gift from Martin Higgs (University of Birmingham). All cell lines were cultured in DMEM plus 10% (v/v) FBS (Sigma-Aldrich) and 1% penicillin/streptomycin in a humidified environment at 37°C with 5% CO_2_. Lentiviral transduced stable cell lines were generated using FuGENE cotransfection of HEK293T with doxycycline-inducible JMJD5 cDNA, constitutively expressing RCCD1 cDNA or doxycycline-inducible HA-JMJD5-P2A-RCCD1-3XFLAG cDNA (details above) plus psPAX2 and pMD2.G packaging vectors. The lentiviral supernatant was filtered and incubated on the recipient cells before selection and growth of successfully transduced cell lines in 1 µg/mL puromycin or 200 µg/mL hygromycin as appropriate.

### Western blotting

Whole cell extracts were prepared in RIPA (25 mM Tris-HCl pH 8, 150 mM NaCl, 0.5% (v/v) sodium deoxycholate (SDS), 1% NP-40) or JIES (100 mM NaCl, 5mM MgCl_2_, 20 mM Tris-HCl pH 7.4, 0.5% (v/v) NP-40) with 1x SIGMAFAST protease inhibitor cocktail (Sigma-Aldrich). Proteins were separated by SDS-PAGE using 12% (v/v) polyacrylamide gels. Samples were subsequently transferred onto PVDF membranes (GE Healthcare) before immunoblotting with the indicated antibodies (Supplemental Figure 17). All antibodies were sourced commercially with the exception of mouse anti-JMJD5 which was a gift from Matsuura Yoshiharu (Osaka University, Osaka, Japan). Chemiluminescent signal was developed using Clarity or Femto-ECL (Bio-Rad) and visualised using a Fusion FX Vilber Imager. Coomassie staining was carried out using Instant Blue (Sigma-Aldrich).

### Immunoprecipitation

Whole cell extracts were prepared in JIES as above. An input fraction was taken and prepared in 6x Laemmli buffer (125 mM Tris-HCl pH 6.8, 6% SDS (w/v), 50% glycerol (v/v), 225 mM dithiothreitol (DTT), 0.1% (w/v) bromophenol blue. Anti-HA agarose (Sigma-Aldrich) or anti-FLAG (Sigma-Aldrich) magnetic beads were added to the remaining whole cell lysates and incubated rotating overnight at 4°C. The beads were then washed six times in JIES before peptide elution using 100 µg FLAG (Generon) or HA (Sigma-Aldrich) peptide in 20 mM Tris-HCl pH8 and 100 mM NaCl. The beads were incubated in elution buffer for 15 minutes at room temperature with shaking at 1400 rpm. The resulting elution was prepared in 6x Laemmli buffer and boiled at 95°C for 5 minutes.

### Mass spectrometry

The HA-JMJD5:RCCD1-3XFLAG complex was anti-FLAG immunopurified from overexpressing HEK293T P2A cells and eluted with 3XFLAG peptide. A control sample of HEK293T cells expressing pTIPZ EV was included to identify non-specific interactors associated with anti-FLAG immunoprecipitation. Samples were extracted, in-solution digested with trypsin, desalted, and analyzed by mass spectrometry. Data were searched against the UniProt Reference Homo sapiens database with PEAKS Studio X Software (Bioinformatics Solutions). Cysteine carbamidomethylation was selected as a ‘fixed’ modification, with oxidation (M) and deamidation (N, Q) as variable modifications. Resulting data was triaged as follows, firstly any proteins identified in the control (EV) samples lacking HA-JMJD5:RCCD1-FLAG were removed. We also removed proteins identified in 50% or more of the screens listed in the ‘Contaminant Repository for Affinity Purification’ (CRAPome) [46]. The resulting top 50 were ranked by percentage coverage and presented in Supplementary Table 4.

### Recombinant JMJD5 protein

pGEX-4T-1 JMJD5 was transformed into BL21(DE3)-competent cells (Promega) and recombinant protein was generated as previously described [13].

### *In vitro* interaction assays

*In vitro* interaction assays were performed by first using anti-FLAG immunoprecipitation as described above. After the wash steps the anti-FLAG magnetic beads were resuspended in 1 mL of interaction buffer (100 mM NaCl and 20 mM Tris pH8). 2.5 µg of recombinant CX3 complex (gift from Steve West, Francis Crick Institute) was added to the samples and incubated at 4°C rotating for 4 hours. Samples were then washed six times in interaction buffer and eluted using 100 µg FLAG peptide as above (Immunoprecipitation).

### *In vitro* hydroxylation assay

As previously described [13].

### Immunofluorescence

Cells were grown on glass coverslips and fixed using 4% (v/v) paraformaldehyde (PFA) at room temperature for 15 minutes. For 53BP1 staining, pre-extraction was performed as previously described [13]. Cells were permeabilised using 0.1% (v/v) Triton X-100 for 10 minutes before blocking in 1% (w/v) BSA (Sigma-Aldrich) in PBS for 1 hour. Coverslips were then stained with the indicated antibodies (Supplemental Figure 17), incubated in DAPI (Invitrogen) for 10 minutes before mounting with ProLong Gold (Cell Signalling). Cells were imaged using a Leica DM6000B.

### DNA fibre assay

Progression of DNA synthesis was monitored using the DNA fibre assay as previously described [13].

### Statistics

Experiments with a minimum of three independent biological repeats were used for statistical analysis and the statistical test used is indicated for each experiment. Graphpad Prism was used to perform 1-way ANOVA and Kruskal-Wallis tests.

### Structural modelling

The interaction between JMJD5 and RCCD1 was predicted using AlphaFold 3 [47]. Predicted local distance difference tests (pLDDT) and interface predicted template modelling (iPTM) was performed within AlphaFold 3 to provide information about the structural accuracy of the generated models. The resulting predictions were analysed using Chimera X.

### Analytical gel filtration

Cells expressing JMJD5-RCCD1 were washed twice with PBS and lysed in a buffer consisting of 20 mM Tris pH 8.0, 100 mM NaCl, 5 mM MgCl_2_, 0.5% (v/v) NP-40, EDTA-free protease inhibitor tablets, and 0.1 % (v/v) turbonuclease. The cell lysate was then centrifuged at 1500 rpm for 5 min, with the supernatant then incubated overnight with 40 μL of Anti-FLAG M2 magnetic beads per 4 mL of cell lysate. Beads were centrifuged at 1500 rpm for 2 min and subsequently washed 6 times in 20 mM Tris pH 8.0, 100 mM NaCl, 5 mM MgCl2, 0.5% (v/v) NP-40. JMJD5-RCCD1 was eluted from the beads using 100 μg/mL FLAG peptide in 20 mM Tris pH 8.0, 100 mM NaCl. The fractions containing JMJD5-RCCD1 were pooled and concentrated before additional purification on a Superdex 200 10/300 gel filtration column (GE Healthcare) equilibrated in 20 mM Tris pH 8.0, 100 mM NaCl, 0.5 mM TCEP. The peak absorbance at 280 nm of JMJD5-RCCD1 was then compared to a set of protein standards of known molecular weights to allow for Mw approximation.

### Multiple sequence alignment

JMJD5 protein sequences used for evolutionary conservation analysis were identified through BLAST searches of NCBI databases. Sequences were aligned in MEGAX using the MUSCLE algorithm [48] and shaded using the Texshade LaTeX package.

### Illustration of JMJD5 mutation plot

Somatic mutation data for JMJD5 was obtained from the COSMIC and cBioPortal databases and imported into R for analysis. Amino acid substitutions and variant classifications were cleaned and reformatted to extract the corresponding residue positions within the encoded protein. Lollipop plot was generated using the *G3viz* package in R. Mutation sites were mapped to a linear protein schematic, with lollipop stems and markers drawn from the processed mutation table. Variant classifications were encoded by colour to distinguish between mutation types. Plot appearance and layout were further refined using *ggplot2*.

## Acknowledgements

T.J.K. was supported by an MRC-DTP award to T.J.K. and M.L.C. S.C.F., E.H. and U.B. were funded by a Cancer Research UK Programme Foundation Award to M.L.C. (C33483/A2567). S.J.S. was supported by a transitional award from the Francis Crick Institute (CR2019/003DK). The work was also supported by an MRC project grant to M.L.C. and S.J.S (APP23111).

## Author contributions

T.J.K. and S.C.F. designed and performed the cell biology experiments, analysed the resulting data, and generated the corresponding figures. T.J.K. led studies into the role of RCCD1 in replication fidelity in tumour cell lines, the JMJD5:RCCD1 interaction and its importance for replication fidelity and replication fork restart, and proteomic screens that identified CX3. S.C.F. led studies into the role of JMJD5 in replication fidelity in tumour cell lines, cancer genetic analyses, JMJD5 activity assays, and replication stress analysis in cells reconstituted with JMJD5 cancer mutants. T.J.K., S.C.F., and M.L.C wrote the manuscript. S.J.S. and G.H. performed structural and biophysical analyses. M.L.C. co-designed the experiments and directed the project with S.J.S., S.C.F., and T.J.K. All authors approved the final version of the manuscript.

