## Supplementary Material for "Inactivation of a protein hydroxylase complex impairs replication fork restart in cancer and neurodevelopmental disorders"

**Supplemental Figure 1**

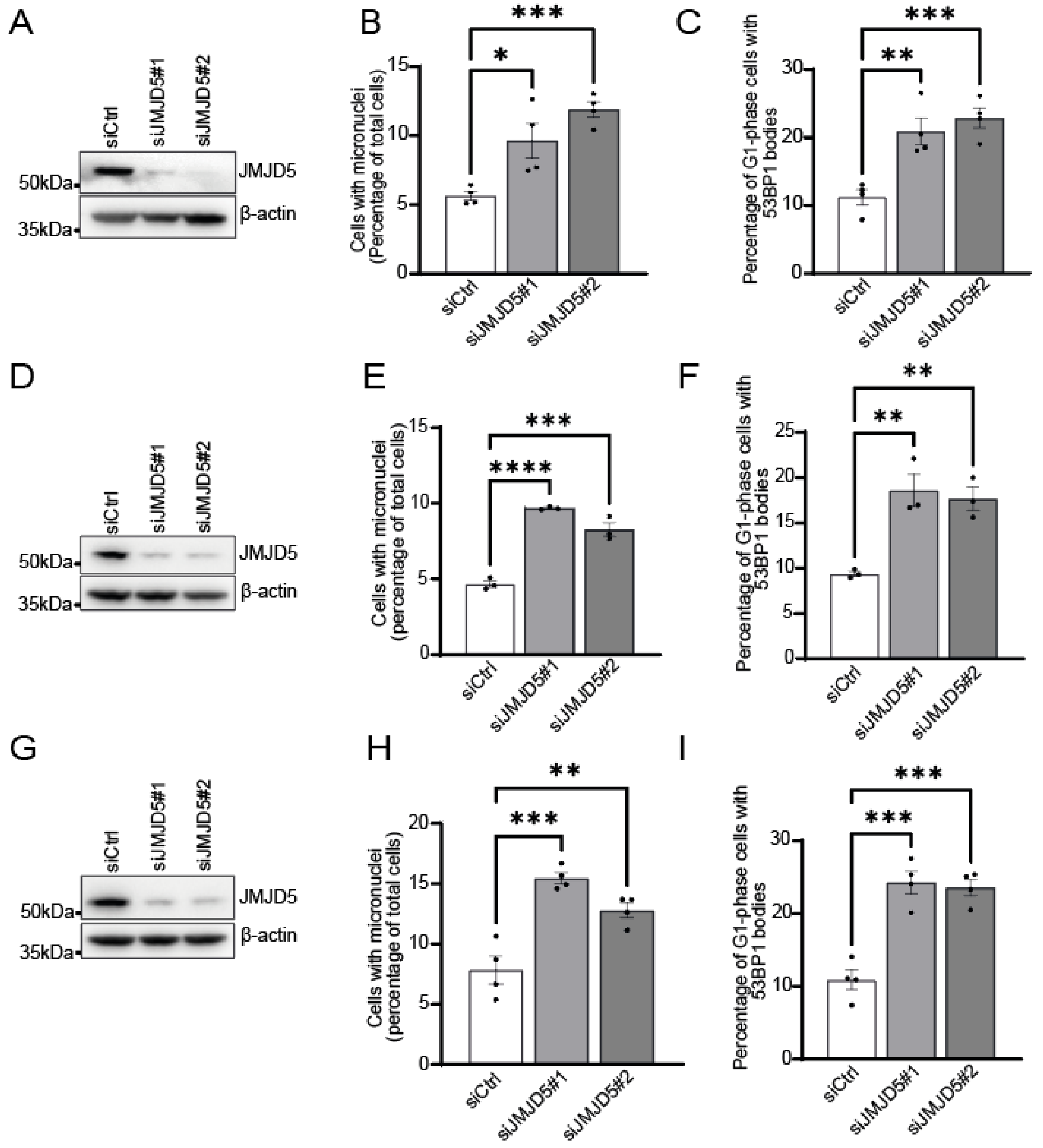


**Supplemental Figure 1. Cancer cell lines U2OS, HeLa and Saos-2 show increased spontaneous 53BP1 bodies in G1 phase and micronuclei following JMJD5 knockdown.** Western blot validation of JMJD5 knockdown using two independent sequences in **(A)** U2OS osteosarcoma cells, **(D)** HeLa **(G)** Saos-2 osteosarcoma cells. JMJD5 knockdown leads to increased micronuclei formation in **(B)** U2OS, **(E)** HeLa and **(H)** Saos-2 cells. Knockdown of JMJD5 also leads to increased formation of 53BP1 bodies in **(C)** U2OS, **(F)** HeLa and **(I)** Saos-2 cells. Data represents mean ± SEM from 3 (**B**, **C**, **E**, **F**) or 4 (**H**, **I**) independent experiments. For 53BP1 bodies and micronuclei a minimum of 250 and 500 cells respectively were counted. Statistical analyses used 1-way ANOVA with Bonferroni's post hoc test. *P≤ 0.05, **P ≤ 0.01, ***P ≤ 0.001, ****P ≤ 0.001.

**Supplemental Figure 2**

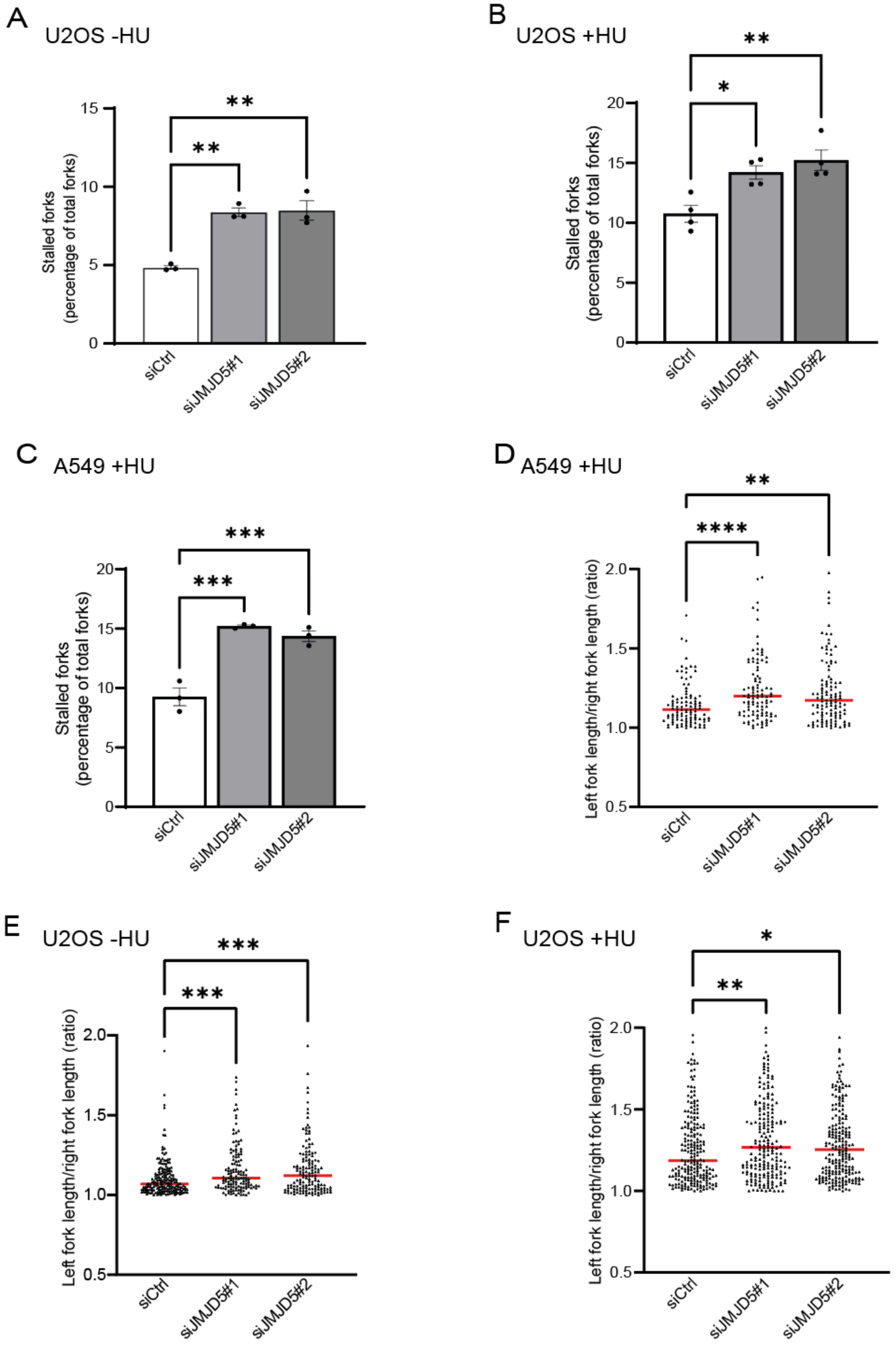

**Supplemental Figure 2. A549 and U2OS cells show increased basal and HU-induced stalled forks and fork asymmetry following JMJD5 knockdown.** U2OS cells treated with two independent siRNA sequences targeting JMJD5 showed significant increases in basal **(A)** and HU-induced **(B)** stalled forks. JMJD5 knockdown in A549 cells treated with HU showed significant increases in **(C)** stalled forks and **(D)** fork asymmetry. Increased fork asymmetry was also observed in U2OS cells in both basal **(E)** and HU treated conditions **(F).** In **(A), (C)** data represent mean ± SEM from 3 independent experiments, 4 independent experiments in **(B)**. In **(D-F)** individual data points from three independent experiments are shown with red line indicating median. For stalled forks a minimum of 250 forks were counted per condition per independent experiment. For fork asymmetry at least 50 structures were measured per sample per independent experiment. Statistical analyses used 1-way ANOVA with Bonferroni's post hoc test **(A-C)** or Kruskal-Wallis with Dunn's correction **(D-F)**. $^{**}P \leq 0.01$, $^{***}P \leq 0.001$, $^{****}P \leq 0.0001$.

**Supplemental Figure 3**

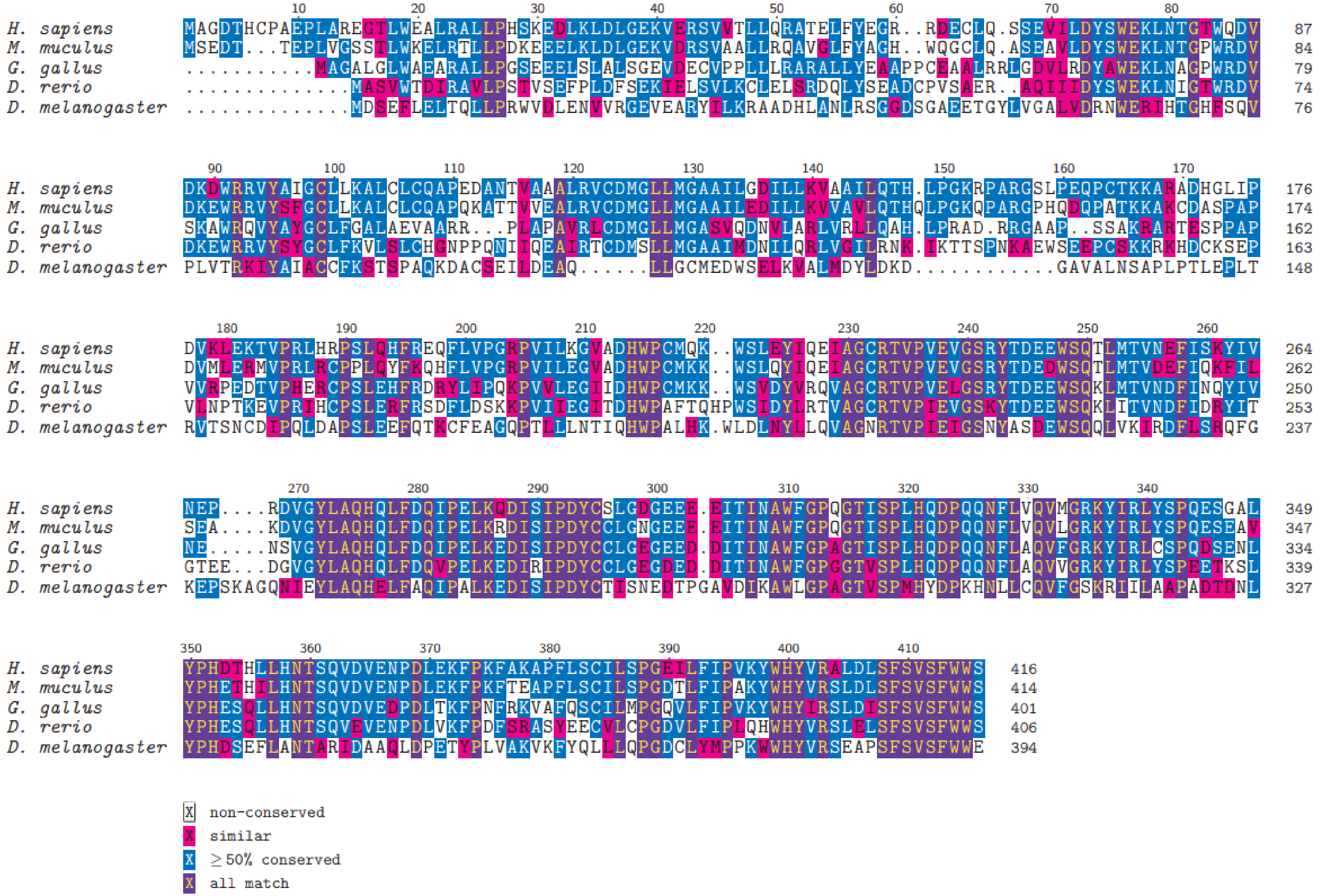

**Supplemental Figure 3. JMJD5 Conservation Analysis.** Multiple sequence alignment (MSA) of JMJD5 generated using MEGA7 and shaded using the Texshade LaTeX package. Shading shows degree of conservation, as indicated at the bottom of the figure.

**Supplemental Figure 4**

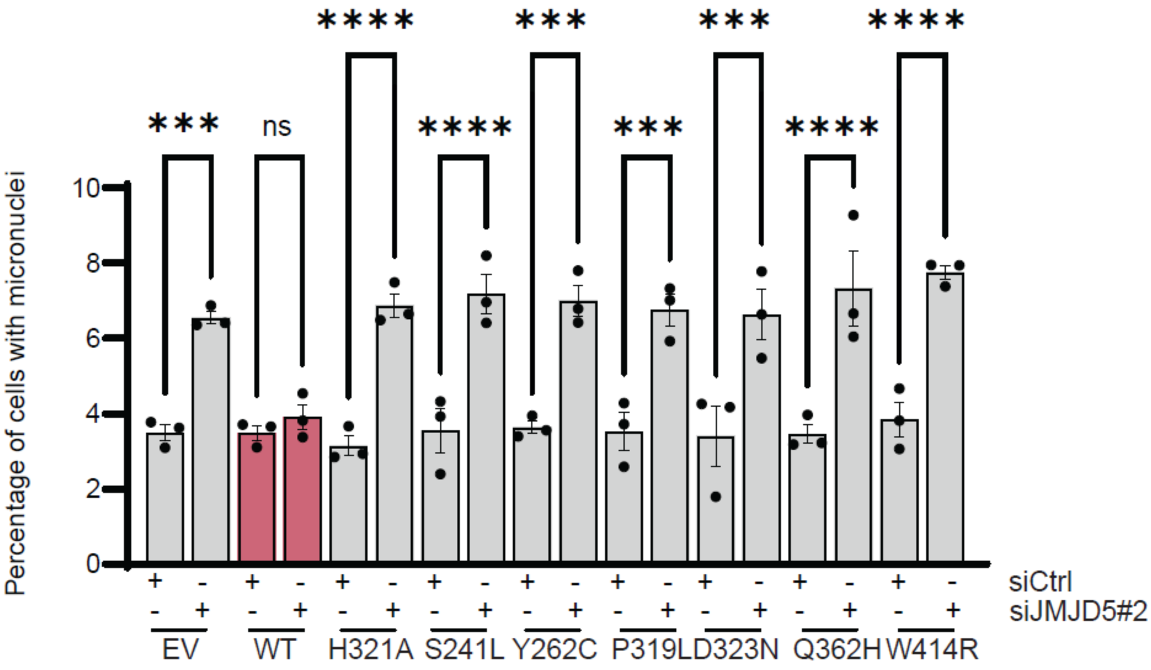


**Supplemental Figure 4. JMJD5 cancer variants are unable to rescue siRNA induced increases in micronuclei.** pTIPZ-FLAG JMJD5 A549 cell lines expressing the indicated JMJD5 cancer variants were treated with 10 ng/mL doxycycline for one week prior to transfection with siCtrl or siJMJD5#2. Cells were fixed after 72 h and micronuclei quantified. Re-expression of WT JMJD5 was able to reduce micronuclei levels in JMJD5 knockdown cells. In contrast, re-expression of either the catalytically dead H321A mutant, or cancer variants shown to affect JMJD5 activity were unable to reduce micronuclei levels. Data represent mean ± SEM from three independent experiments. A minimum of 500 cells were counted per sample in each experiment. Statistical analysis used 1-way ANOVA with Bonferroni's post hoc test; ns not significant, ***P ≤ 0.001, ****P ≤ 0.0001.

**Supplemental Figure 5**

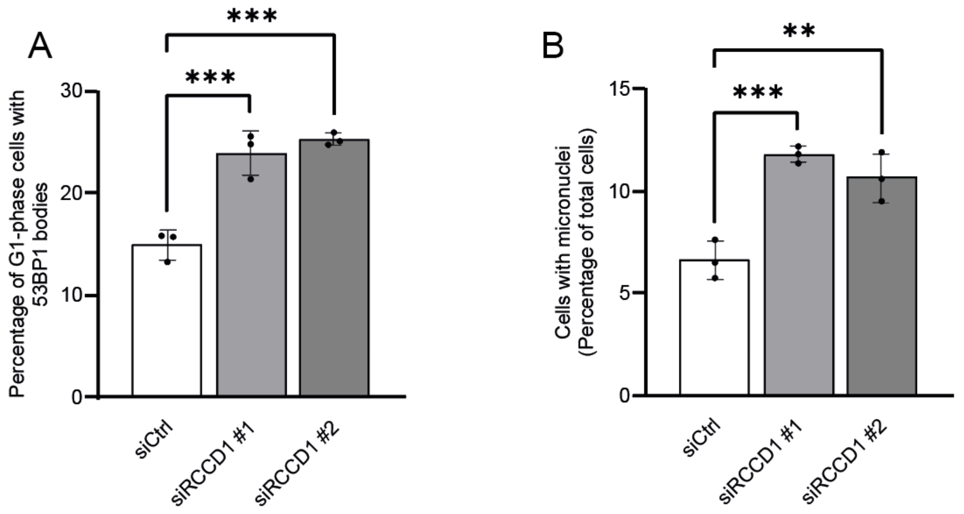


**Supplemental Figure 5. U2OS osteosarcoma cells display increased spontaneous 53BP1 bodies in G1 phase and micronuclei.** Basal levels of **(A)** 53BP1 bodies in G1 phase and **(B)** micronuclei were significantly increased in U2OS cells following knockdown of RCCD1 using two independent siRNAs. Data represent mean ± SEM from 3 independent experiments. For 53BP1 bodies and micronuclei a minimum of 300 and 500 cells respectively were counted. Statistical analyses used 1-way ANOVA with Bonferroni's post hoc test. $^{**}P \leq 0.01$, $^{***}P \leq 0.001$.

**Supplemental Figure 6**

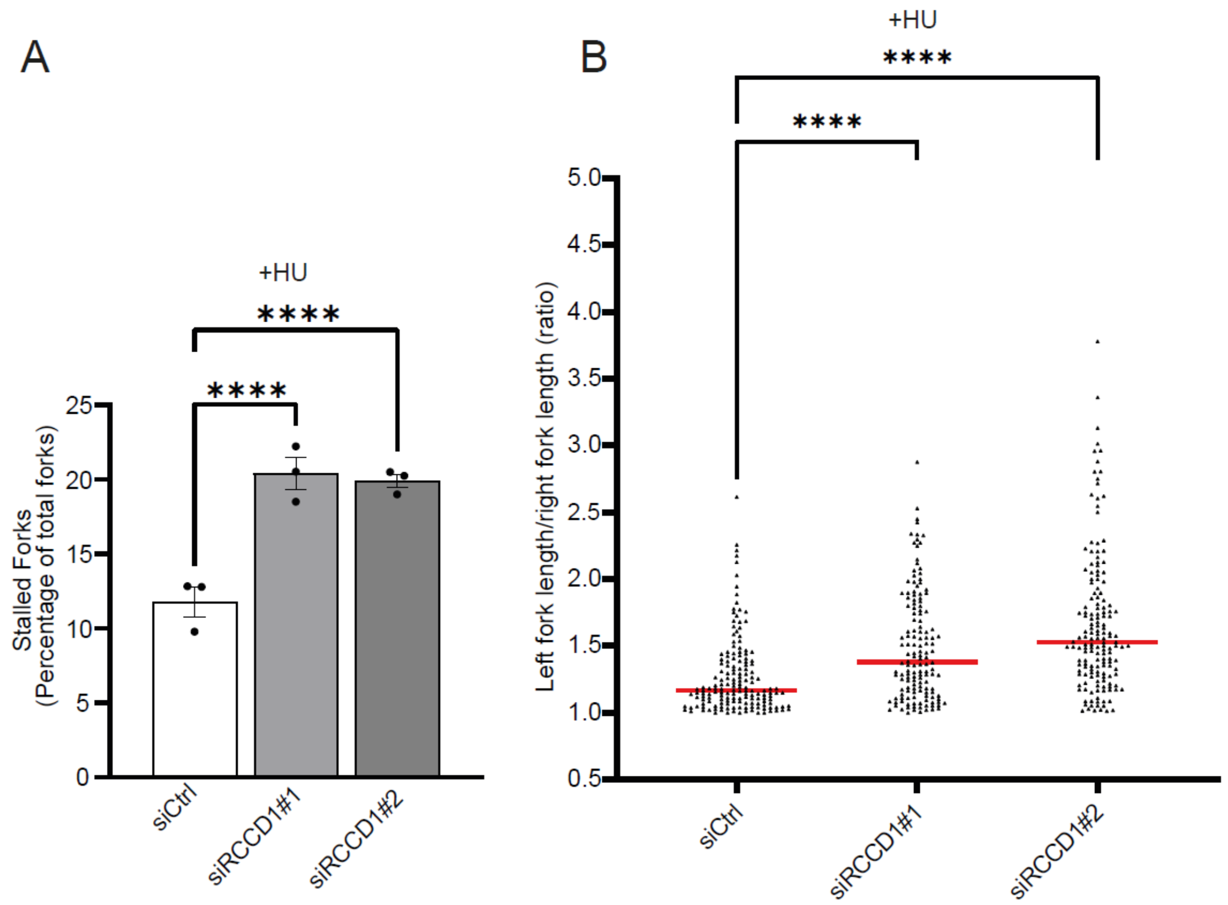


**Supplemental Figure 6. Increased stalled forks and fork asymmetry following RCCD1 knockdown in A549 cells is exacerbated by HU treatment.** Significant increases in **(A)** stalled forks and **(B)** fork asymmetry was observed in HU treated A549 cells following RCCD1 knockdown. Data represent mean ± SEM from 3 independent experiments. For stalled forks a minimum of 250 forks were counted per condition per independent experiment. For fork asymmetry at least 50 structures were measured per sample per independent experiment. Statistical analyses used 1-way ANOVA with Bonferroni's post hoc test **(A)** or Kruskal-Wallis with Dunn's correction **(B)**. ****$P \leq 0.0001$.

**Supplemental Figure 7**

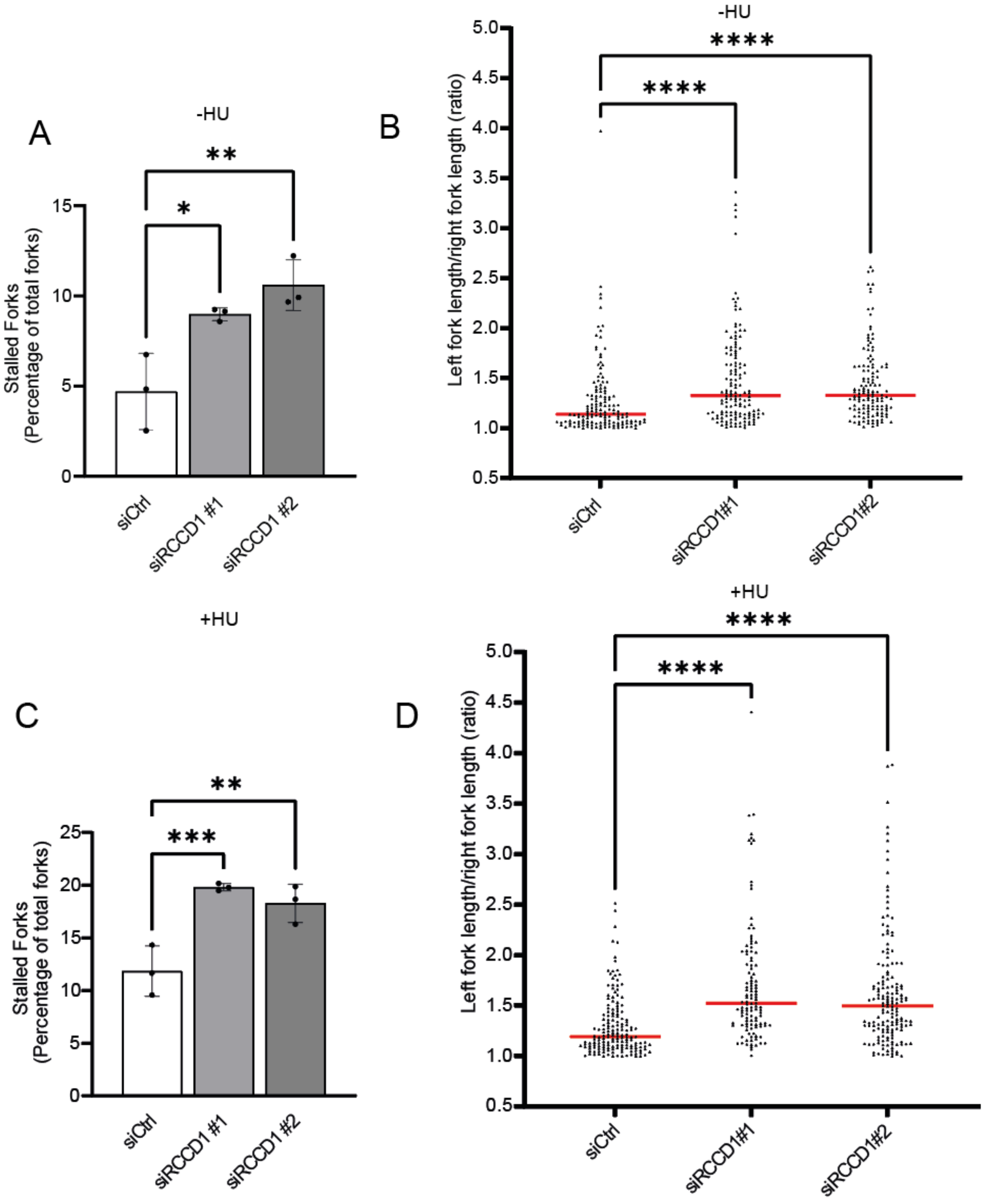


**Supplemental Figure 7. U2OS cells display increased basal and HU-induced stalled forks and fork asymmetry following RCCD1 knockdown.** U2OS cells treated with siRNA against RCCD1 showed significant increases in basal levels of (A) stalled forks and (B) fork asymmetry. With both (C) stalled forks and (D) fork asymmetry exacerbated by treatment with HU. Data represent mean ± SEM from 3 independent experiments. For stalled forks a minimum of 250 forks were counted per condition per independent experiment. For fork asymmetry at least 50 structures were measured per sample per independent experiment. Statistical analyses used 1-way ANOVA with Bonferroni's post hoc test **(A and C)** or Kruskal-Wallis with Dunn's correction **(B and D)**. *$P \leq 0.05$, **$P \leq 0.01$, ***$P \leq 0.001$, ****$P \leq 0.0001$.

**Supplemental Figure 8**

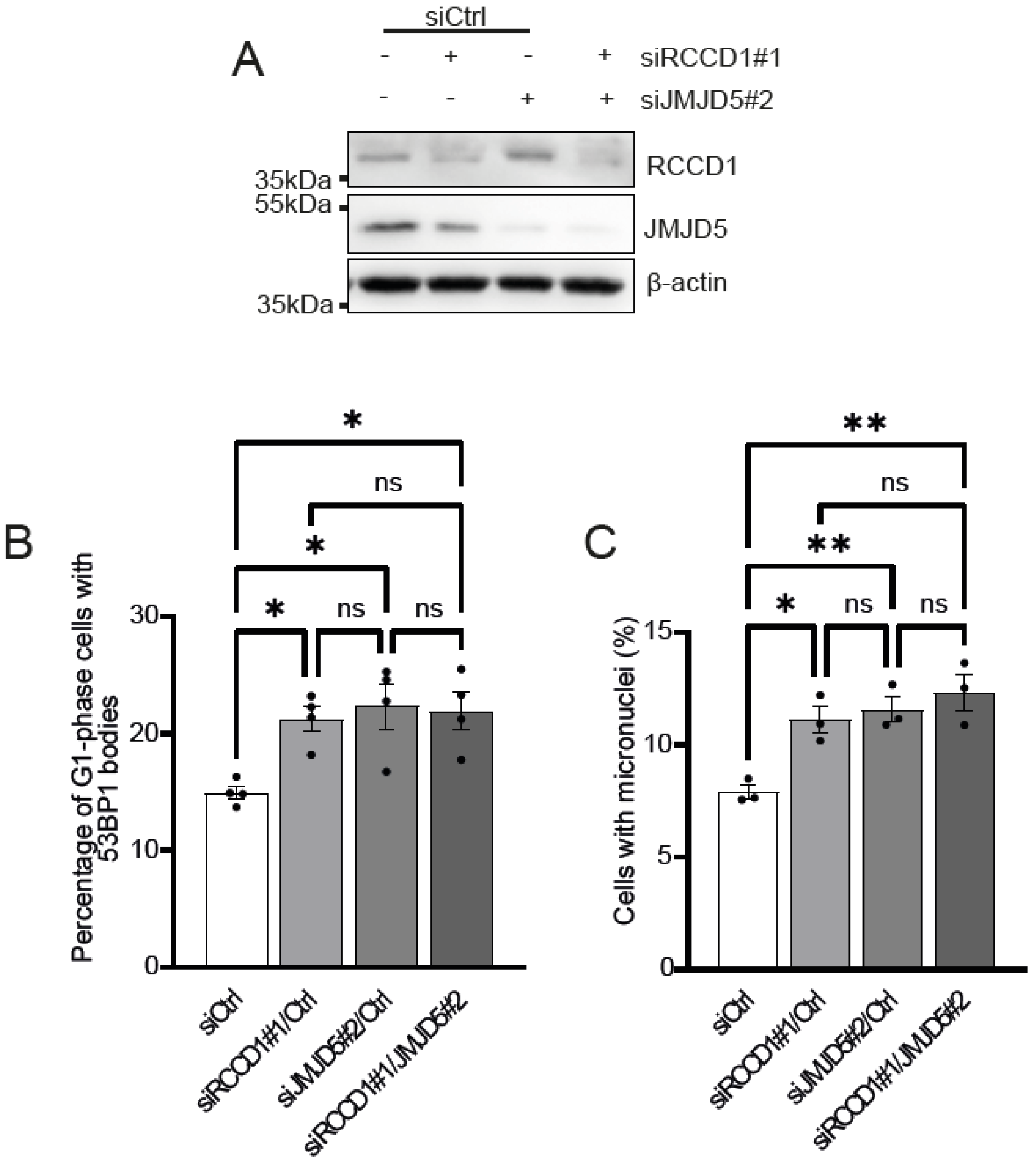


**Supplemental Figure 8. Increased replication stress phenotype associated with JMJD5 and RCCD1 knockdown in U2OS cells is epistatic.** U2OS cells treated with the indicated siRNA combinations with successful knockdown confirmed via **(A)** western blot, which displayed significant increases in basal levels of **(B)** 53BP1 bodies and **(C)** micronuclei following JMJD5/RCCD1 knockdown, which was not exacerbated by co-depletion. Data represent mean ± SEM from 4 **(B)** or 3 **(C)** independent experiments. For 53BP1 bodies and micronuclei a minimum of 300 and 500 cells respectively were counted. Statistical analyses used 1-way ANOVA with Bonferroni's post hoc test. $^{*}P \leq 0.05$, $^{**}P \leq 0.01$; ns = non-significant.

**Supplemental Figure 9**

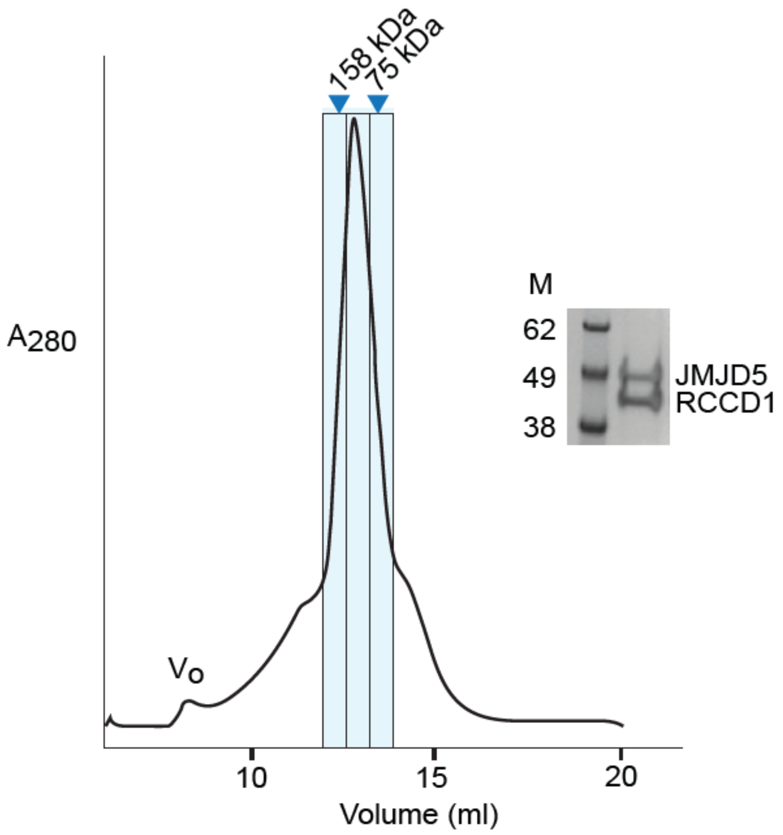


**Supplemental Figure 9. Analytical Gel filtration chromatography of recombinant JMJD5:RCCD1 isolated from HA-JMJD5-P2A-RCCD1-3XFLAG HEK293T cells.** Gel filtration was carried out on a Superdex S-200 Increase10/300 column. Coomassie gel is of the fractions indicated by the blue markers. The blue arrows indicated the calibration molecular weight markers.

**Supplemental Figure 10**

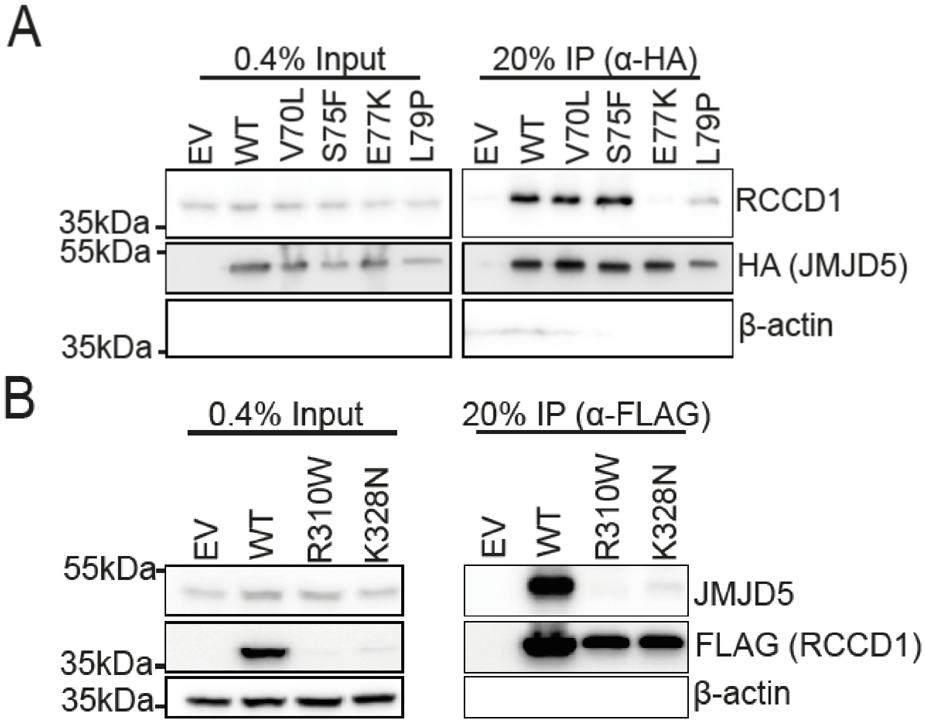


**Supplemental Figure 10 RCCD1 cancer variants affect RCCD1 expression and JMJD5 interaction (A)** JMJD5 E77K completely ablates interaction with RCCD1. HEK293T cells were transfected with control pEF6 (EV), pEF6-HA-JMJD5(WT) or selected cancer mutations pEF6-HA-JMJD5(V70L, S75F, E77K or L79P) followed by anti HA-immunoprecipitation and western blotting with the indicated antibodies. **(B)** Selected cancer mutations R310W and K328N reduced RCCD1 protein expression and JMJD5 binding. Lysates from HEK293T cells transfected with control pIHZ (EV), pIHZ-RCCD1-FLAG(WT) or selected cancer mutations pIHZ-RCCD1-FLAG (R310W or K328N) were anti-FLAG immunoprecipitated and western blotted with the indicated antibodies.

**Supplemental Figure 11**

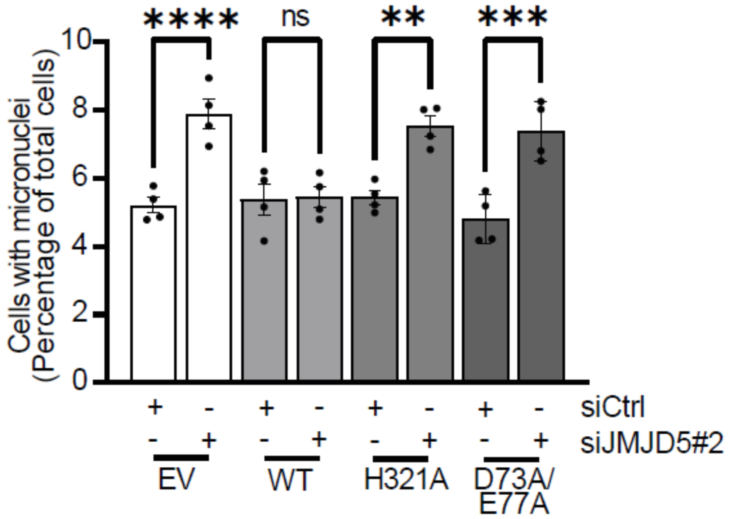


**Supplemental Figure 11. JMJD5 catalytic activity and RCCD1 binding are required to maintain genome stability in A549 cells.** Reconstitution of JMJD5-WT but not catalytically dead (H321A) or the RCCD1 binding mutant (D73A/E77A) was able to rescue basal increase in micronuclei. Data represents mean ± SEM for 4 independent experiments. For micronuclei, a minimum of 500 cells were counted per sample. Statistical analyses used 1-way ANOVA with Bonferroni’s post hoc test **(A).** **P ≤ 0.01, ***P ≤ 0.001, ****P ≤ 0.0001. ns = non-significant.

**Supplemental Figure 12**

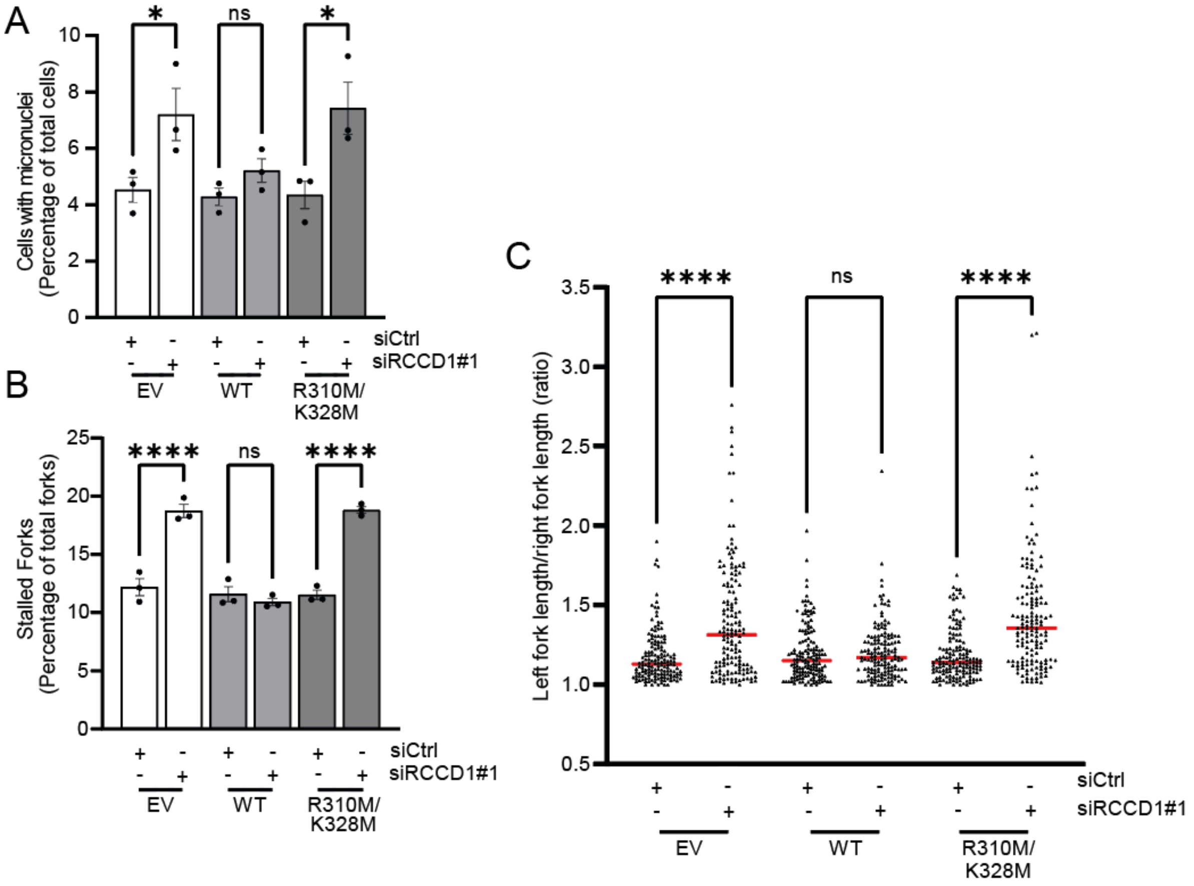


**Supplemental Figure 12. RCCD1 requires JMJD5 binding to maintain replication fidelity in A549 cells.** Reconstitution of WT RCCD1 but not the JMJD5 binding mutant (R310M/K328M) was able to rescue **(A)** basal increase in micronuclei and **(B)** stalled forks and **(C)** fork asymmetry. Data represent mean ± SEM from 3 independent experiments. For micronuclei, a minimum of 500 cells were counted per sample. For stalled forks a minimum of 250 forks were counted per condition per independent experiment. For fork asymmetry at least 50 structures were measured per sample per independent experiment. Statistical analyses used 1-way ANOVA with Bonferroni's post hoc test **(A** and **B)** or Kruskal-Wallis with Dunn's correction **(C)**. *P ≤ 0.05, ****P ≤ 0.0001. ns = non-significant.

**Supplemental Figure 13**

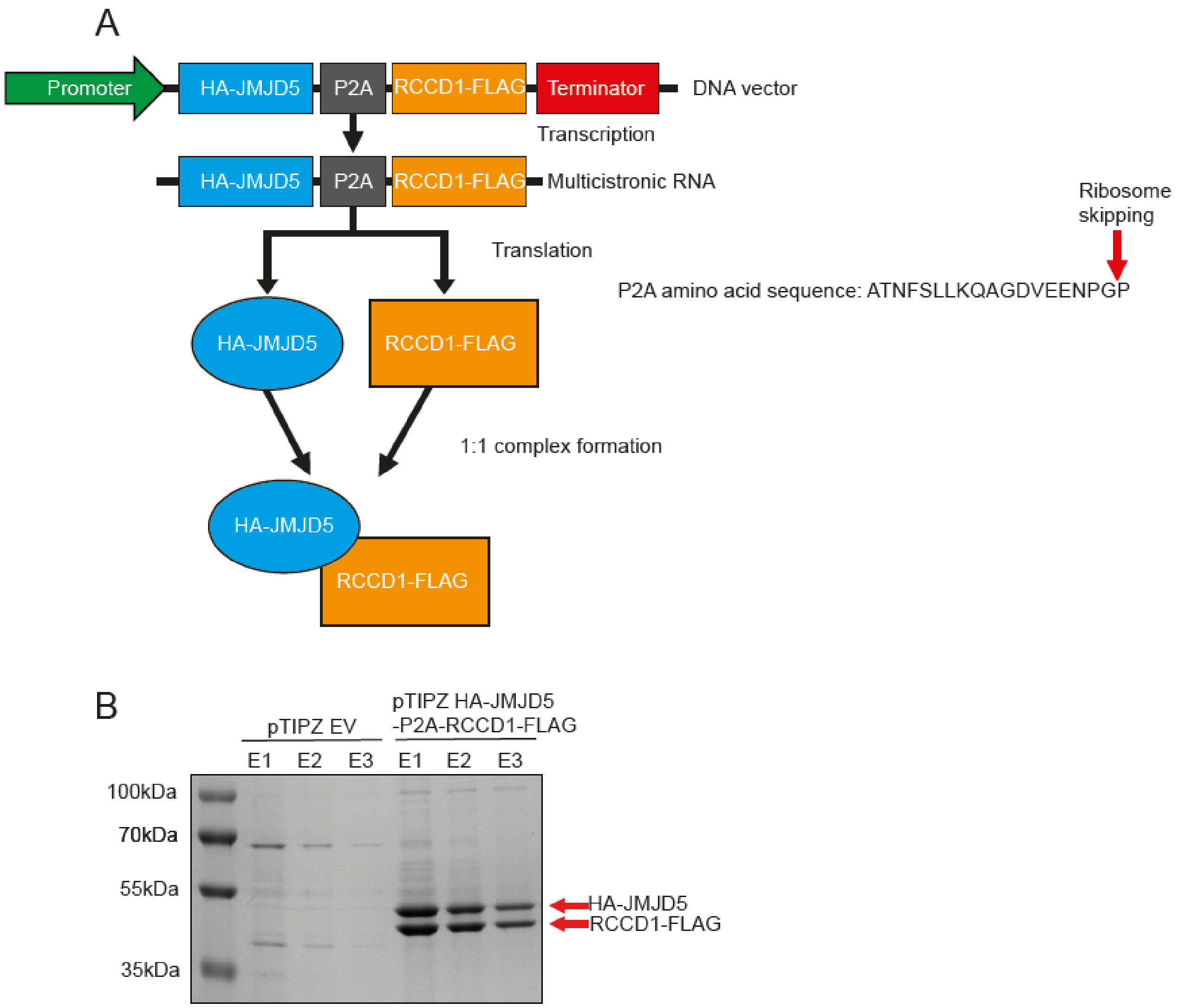


**Supplemental Figure 13. Developmental and validation of JMJD5/RCCD1 co-overexpression system. (A)** Schematic of HA-JMJD5-P2A-RCCD1-FLAG co-overexpression system. cDNA vector is transcribed into a multicistronic mRNA, during translation the porcine-adenovirus sequence causes ribosome skipping between the final glycine and proline, translating two separate proteins from a single mRNA. **(B)** Validation of JMJD5/RCCD1 1:1 heterodimeric complex formation from P2A co-overexpression system. Lysates from control (pTIPZ EV) or pTIPZ HA-JMJD5-P2A-RCCD1-FLAG HEK293T cells were subjected to anti-FLAG immunoprecipitation with three subsequent elutions (E1, E2 and E3) followed by SDS-PAGE and Coomassie staining.

**Supplemental Figure 14**

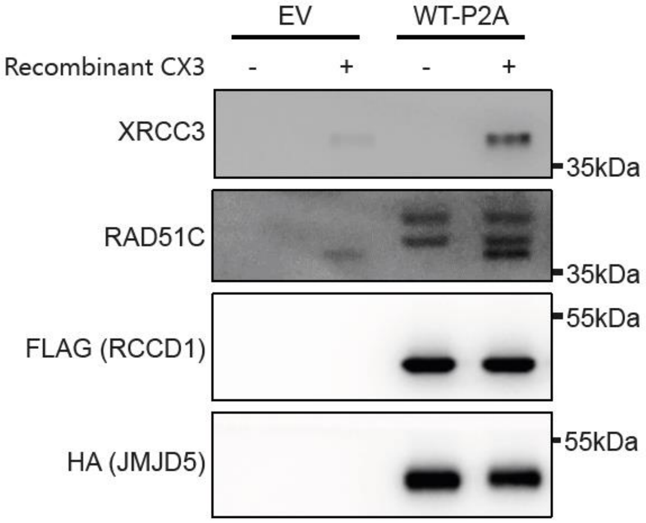


**Supplemental Figure 14. The JMJD5:RCCD1 complex interacts with the CX3 complex of RAD51 paralogs *in vitro.*** Recombinant JMJD5:RCCD1 protein was isolated from HEK293T cells expressing EV or WT JMJD5:RCCD1 protein complex followed by incubation with recombinant CX3 (gift from Steve West). The resulting samples were western blotted with the indicated antibodies.

**Supplemental Figure 15**

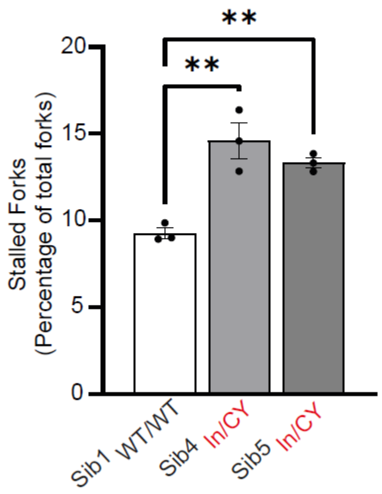


**Supplemental Figure 15. Primary patient fibroblasts from affected JMJD5 patients display increased deficiency in replication fork restart.** Affected patient primary cells showed significantly increased replication fork restart defects compared to non-affected siblings. Statistical analysis used 1-way ANOVA with Bonferroni's post hoc test. $^{**}P \le 0.01$, ns = non-significant.

**Supplemental Figure 16**

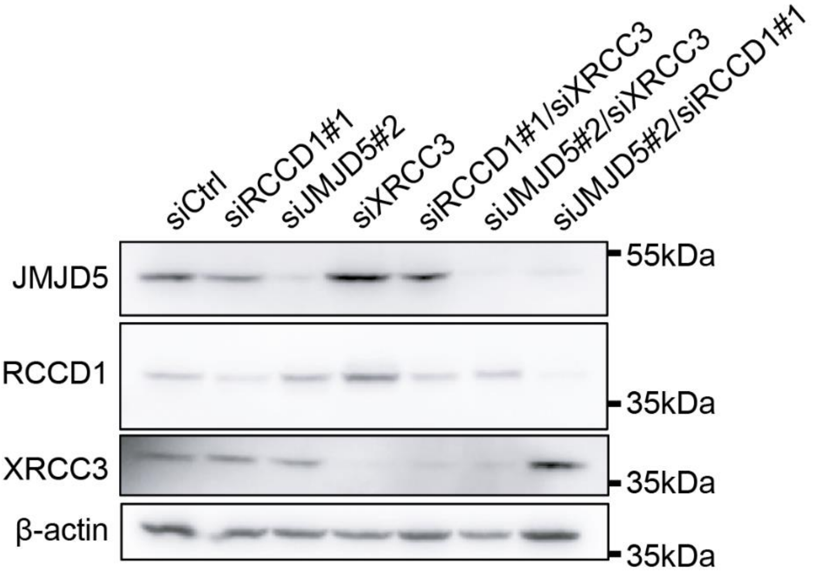


**Supplemental Figure 16. Western blot validation of successful siRNA knockdown of JMJD5, RCCD1 and XRCC3 epistasis experiments.** A549 cells were treated with the indicated siRNA combinations and successful knockdown confirmed via western blot.

**Supplemental Figure 17**

Western blotting

| Target | Antibody |
|---|---|
| α-Rabbit-HRP | Cell Signalling Technology 7074S |
| α-Mouse-HRP | Cell Signalling Technology 7076S |
| β-Actin | HRP-tagged, Abcam ab49900 |
| FLAG | HRP-tagged Sigma-Aldrich, A8592 |
| HA | HRP-tagged Sigma-Aldrich, (12013819001) |
| JMJD5 | Matsuura Yoshiharu Laboratory, Japan |
| RAD51 | Abcam ab63801 |
| RAD51B | Santa Cruz sc-377192 |
| RAD51C | Novus Biologicals NB100-177 |
| RCCD1 | Abcam ab122570 |
| RPA | Calbiochem NA18 |
| XRCC3 | Novus Biologicals NB100-165SS |

Immunofluorescence

| Target | Antibody |
|---|---|
| CENPF | BD Biosciences, 610768 |
| FLAG | Sigma-Aldrich, F1804 |
| 53BP1 | Bio-Techne, NB100-904V |
| BrdU (recognises CldU) | Abcam, ab6326 |
| BrdU (recognises IdU) | BD Biosciences, 347580 |
| Mouse 555nm | Thermofisher, A31570 |
| Mouse 488nm | Thermofisher, A10684 |
| Rabbit 488nm | Thermofisher, A11070 |
| Rat 555nm | Thermofisher, A-21434 |

siRNA sequences

| Sequence name | Catalogue Number |
|---|---|
| siJMJD5#1 | SASI_Hs01_00082889 |
| siJMJD5#2 | SASI_Hs01_00082891 |
| siRCCD1#1 | SASI_Hs01_00084850 |
| siRCCD1#2 | SASI_Hs01_0084852 |
| siXRCC3 | SASI_Hs01_00321675 |
